# aMeta: an accurate and memory-efficient ancient Metagenomic profiling workflow

**DOI:** 10.1101/2022.10.03.510579

**Authors:** Zoé Pochon, Nora Bergfeldt, Emrah Kırdök, Mário Vicente, Thijessen Naidoo, Tom van der Valk, N. Ezgi Altınışık, Maja Krzewińska, Love Dalen, Anders Götherström, Claudio Mirabello, Per Unneberg, Nikolay Oskolkov

**Author notes:** shared authorship (Zoé Pochon, Nora Bergfeldt, Anders Götherström, Claudio Mirabello, Per Unneberg, Nikolay Oskolkov).

## Abstract

Analysis of microbial data from archaeological samples is a rapidly growing field with a great potential for understanding ancient environments, lifestyles and disease spread in the past. However, high error rates have been a long-standing challenge in ancient metagenomics analysis. This is also complicated by a limited choice of ancient microbiome specific computational frameworks that meet the growing computational demands of the field. Here, we propose aMeta, an accurate ancient Metagenomic profiling workflow designed primarily to minimize the amount of false discoveries and computer memory requirements. Using simulated ancient metagenomic samples, we benchmark aMeta against a current state-of-the-art workflow, and demonstrate its superior sensitivity and specificity in both microbial detection and authentication, as well as substantially lower usage of computer memory. aMeta is implemented as a Snakemake workflow to facilitate use and reproducibility.

## Introduction

Historically, ancient DNA (aDNA) studies have focused on human and faunal evolution, extracting and analyzing predominantly eukaryotic aDNA [1–3]. With the development of Next Generation Sequencing (NGS) technologies, it was demonstrated that endogenous microbial communities aDNA from eukaryotic remains, which was previously treated as a sequencing by-product, can provide valuable information about ancient pandemics, lifestyle and population migrations in the past [4–6]. Modern technologies have made it possible to study not only ancient microbiomes populating eukaryotic hosts, but also sedimentary ancient DNA (sedaDNA), which has rapidly become an independent branch of palaeogenetics, delivering unprecedented information about hominin and animal evolution without the need to analyze historical bones and teeth [7–12]. Previously available in microbial ecology, meta-barcoding methods lack validation and authentication power, and therefore shotgun metagenomics has become the *de facto* standard in ancient microbiome research [13]. However, accurate detection, abundance quantification and authentication analysis of microbial organisms in ancient metagenomic samples remains challenging [14]. This is related to the limited amount of microbial aDNA and the exceptional variety of both endogenous and invasive microbial communities that have been populating ancient samples when living and post-mortem. In particular, the presence of modern contamination can introduce biases in the analysis of aDNA data. All of these technical and biological factors can lead to a high rate of false-positive and false-negative microbial identifications in ancient metagenomic studies [15].

When screening for the presence of microbial organisms with available reference genomes, we aim to assign a taxonomic label for each aDNA sequence. For this purpose, two dominant approaches are: composition, aka *k*-mer taxonomic classification, and alignment based methods. For the former, the Kraken family of tools [16, 17] is one of the most popular in ancient metagenomics, while for the latter, general purpose aligners such as BWA [18] and Bowtie2 [19], as well as MALT [20], which was specifically designed for the analysis of metagenomic data, are among the most commonly used.

Unlike the alignment approach, where each aDNA sequence is positioned along the reference genome based on its similarity to it, the *k*-mer taxonomic classification uses a lookup database containing *k*-mers and Lowest Common Ancestor (LCA) information for all organisms with available reference genomes. DNA sequences are classified by searching the database for each *k*-mer in a sequence, and then using the LCA information to determine the most specific taxonomic level for the sequence. Advantages of the classification-based approach are high speed and a wide range of candidates (database size), while disadvantages include a difficulty in validation and authentication that can often lead to a high error rate of the classification-based approach. In contrast, the alignment-based approach with e.g. MALT provides more means of validation and authentication, while being relatively slow, more resource-demanding and heavily dependent on the selection of reference sequences included in the database. Technical limitations such as computer memory (RAM) often hinder the inclusion of a large amount of reference sequences into the database which might result in a high false-negative rate of microbial detection. In practice, due to the very different nature of the analyses and reference databases used, the outputs from classification and alignment approaches often contradict each other, bringing additional confusion to the ancient metagenomics research community. In fact, both approaches have strengths but also profound weaknesses that can lead to substantial analysis error, if not properly taken into account.

Here, we define two types of errors common to ancient metagenomics: 1) the detection error, and 2) the authentication error. The detection error comes from a difficulty to correctly identify microbial presence or absence irrespective of the ancient status. This can happen due to many reasons such as overly relaxed or too conservative filtering. This error is not specific to ancient metagenomics but represents a general challenge that is also valid for the field of modern metagenomics. In contrast, the authentication error is specific to ancient metagenomics and caused by modern contamination that is typically present in archaeological samples. Often, inaccurate data processing and handling can lead to the erroneous discovery of a modern contaminant as being of ancient origin, and *vice versa*, of an ancient microbe as being modern. Therefore, the major goals of an ancient microbiome reconstruction are to establish accurate evidence that a microbe a) truly exists in a sample, and b) is of ancient origin.

In this study, we aim to combine the strengths of both classification- and alignment-based approaches to develop an ancient metagenomics profiling workflow, aMeta, with low detection and authentication errors. For this purpose, we use KrakenUniq [21, 22] — which is suitable for working in low-memory computational environments — for initial taxonomic profiling of metagenomic samples and informing MALT reference database construction, followed by LCA-based MALT alignments, and a comprehensive validation and authentication analysis based on the alignments. We report that a KrakenUniq-based selection of microbial candidates for inclusion in the MALT database, dramatically reduces resource usage of aMeta compared to metagenomic profiling with MALT alone. We evaluated our workflow using simulated ancient metagenomic data, and benchmarked it against Heuristic Operations for Pathogen Screening (HOPS) [23], which is probably the most popular and *de facto* standard ancient metagenomic pipeline currently. We demonstrate that due to its additional breadth / evenness of coverage filtering, superior database size, and flexible authentication score system, the combination of KrakenUniq and MALT implemented in our workflow results in a higher sensitivity vs. specificity balance for detection and authentication of ancient microbes compared to HOPS. Importantly, aMeta consumed nearly half as much computer memory as HOPS on a benchmark simulated ancient metagenomic dataset.

## Method

The aMeta workflow overview is shown in Figure 1. It represents an end-to-end processing and analysis framework implemented in Snakemake [24] that accepts raw data as a set of files, usually belonging to a common project, and outputs a ranked list of detected ancient microbial species together with their abundances for each sample, as well as a number of validation and authentication plots for each identified microorganism in each sample. In other words, the workflow leverages a convenient high-level summary of several authentication and validation metrics that evaluate detected microbes based on the evidence of their presence and ancient status.

**Figure 1.**
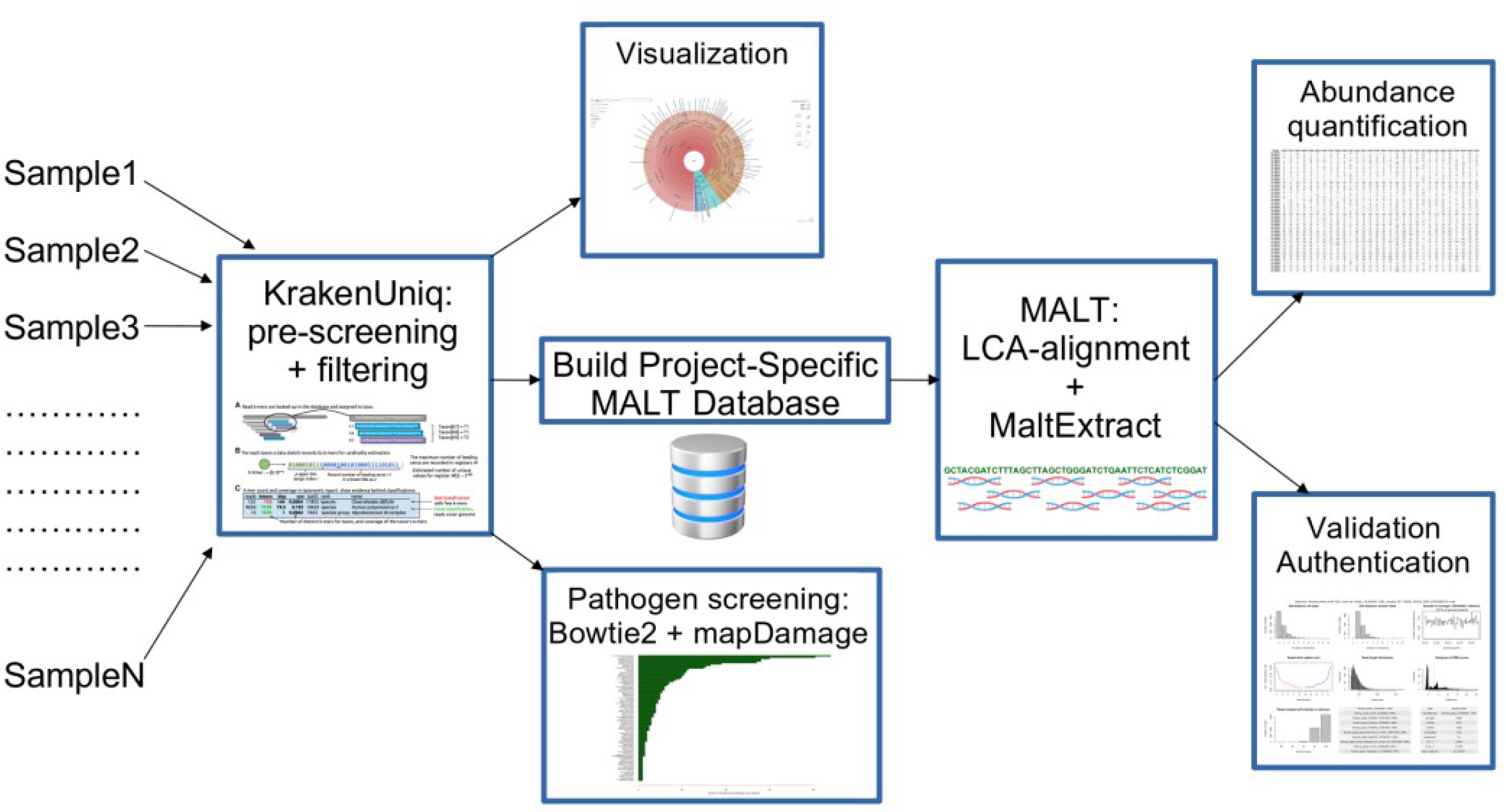
aMeta: ancient metagenomics profiling workflow overview. The workflow represents a combination of taxonomic classification + filtering steps with KrakenUniq that allows to establish a list of microbial candidates for further building a MALT database, running LCA-based alignments with MALT against the database, and performing validation + authentication analysis based on the alignments.

Below we provide a detailed description of each step implemented in aMeta. The workflow accepts raw metagenomic data in a standard *fastq* format, removes sequencing adapters with Cutadapt [25], and selects reads of length above 30 bp which have a good taxonomic specificity. Next, the workflow runs KrakenUniq [21, 22] (below we refer to this step as “pre-screening”), a fast and accurate *k*-mer-based tool which is capable of operating in low-memory computational environments [22]. KrakenUniq performs a taxonomic classification of aDNA sequences, and reports a number of *k*-mers unique to each taxa, which can be considered equivalent to the breadth of coverage information. The number of unique *k-*mers is an essential filter of aMeta which significantly improves its accuracy, see below. Generally, breadth of coverage information is obtained through alignments, therefore the advantage of KrakenUniq is that it is capable of delivering a breadth of coverage estimation via classification without performing explicit alignments.

Figure 2 schematically demonstrates why detecting microbial organisms solely based on depth of coverage (or simply coverage), which is largely equivalent to the number of mapped reads, might lead to false-positive identifications. Suppose we have a toy reference genome of length 4 * *L* and 4 reads of length *L* mapping to the reference genome. When a microbe is truly detected, the reads should map evenly across the reference genome, see Figure 2B. In contrast, in case of misaligned reads, i.e. when reads originating from species A map to the reference genome of species B, it is common to observe “piles’’ of reads aligned to a few conserved regions of the reference genome, which is the case in Figure 2A (see also Supplementary Figure 1 for a real data example, where reads from unknown microbial organisms are forced to map to *Yersinia pestis* reference genome alone). Therefore, we consider the breadth of coverage information delivered by KrakenUniq to be of crucial importance for robust filtering in our workflow.

**Figure 2.**
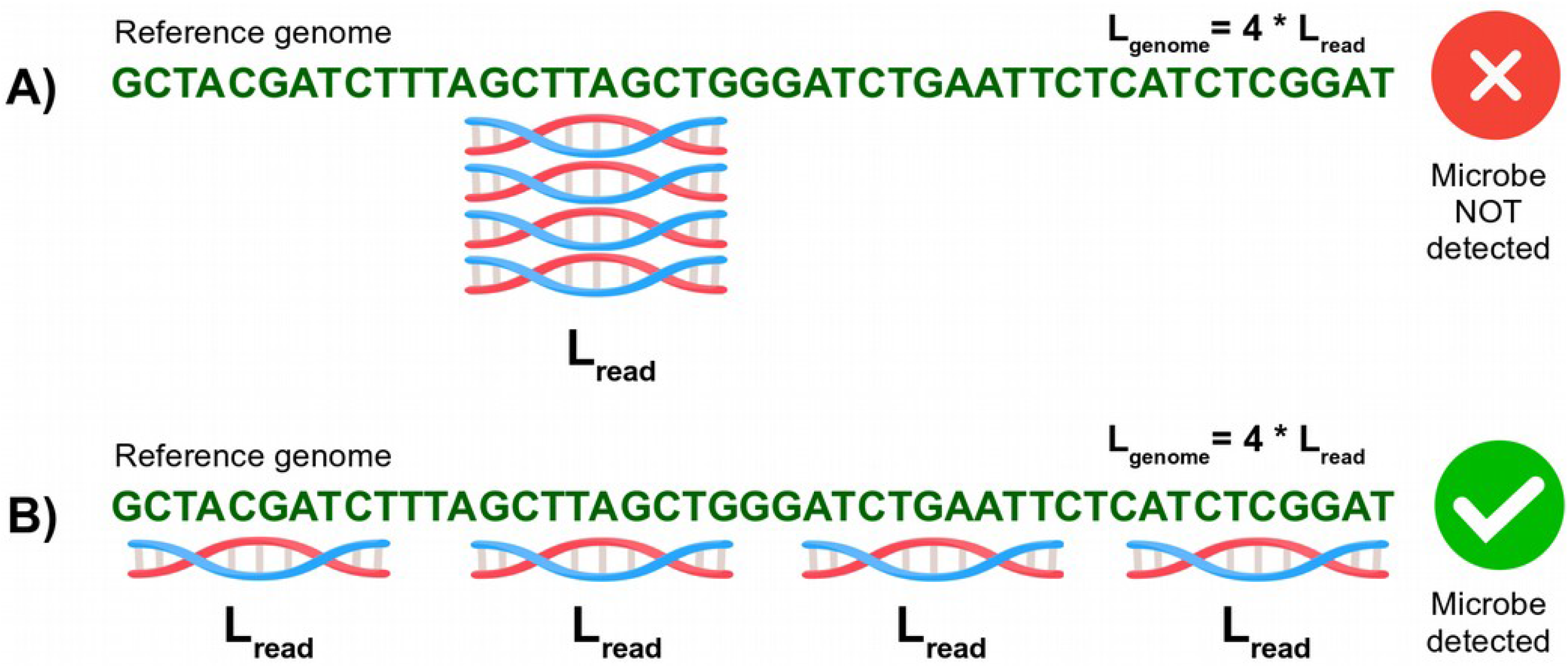
Demonstration of the difference between depth and breadth of coverage concepts. Two read alignment scenarios, A) and B), have identical depth of coverage (or simply coverage) of N_reads_* L_read_ / L_genome_= 4 * L_read_ / 4 * L_read_ = 1X. However, the reads are spread unevenly in case A) and evenly in case B). The latter has a higher breadth of coverage and corresponds to a true-positive hit, while the former, A), scenario is typical for a false-positive microbial detection.

In addition to the filtering with respect to breadth of coverage, low-abundance microbes are removed in aMeta based on their depth of coverage, which is related to the number of reads assigned to each taxa. Filtering by depth of coverage is also important for subsequent validation and authentication steps, as some of these may not be statistically robust enough when performed on low abundant microbes. Therefore, aMeta uses a rather conservative approach, and concentrates on reasonably abundant species with a uniform coverage which are more likely to be truly present in the samples, Figure 3.

**Figure 3.**
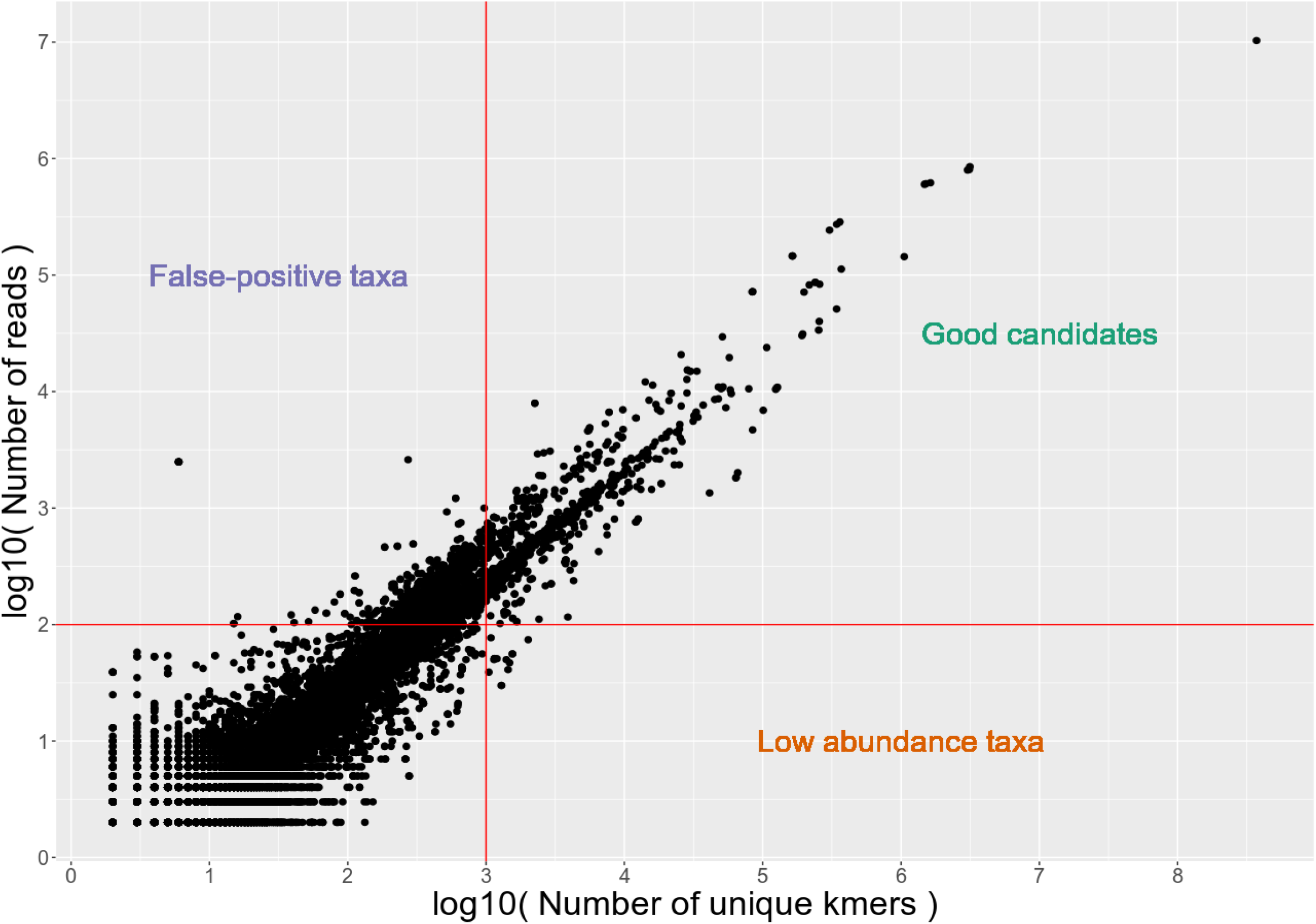
Depth (number of reads) and breadth (number of unique *k*-mers) of coverage filters applied by KrakenUniq to microbial taxa in a metagenomic sample. Taxa with large amounts of non-uniformly mapped reads, and hence low breadth of coverage, are considered to be false-positive identifications (left upper corner). Red solid horizontal and vertical lines mark possible depth (~ 100 reads) and breadth (~1000 unique kmers) of coverage filters applied to KrakenUniq output.

For pre-screening with KrakenUniq, we built two different databases of reference sequences: 1) a full NCBI non-redundant NT database, currently used by default in BLASTN [26], that included all eukaryotic and prokaryotic genomes available at NCBI; 2) a microbial version of NCBI NT database, consisting of all microbial genomes (bacteria, viruses, archaea, fungi, protozoa and parasitic worms) as well as the human genome and only complete eukaryotic genomes from NCBI. The former database can be used for comprehensive screening for both eukaryotic (mammals, plants etc.) and microbial organisms, while the latter is less than half in size and can suffice when performing only microbial profiling. Both databases are publicly available to the wider scientific community through the SciLifeLab Figshare at https://doi.org/10.17044/scilifelab.20205504 and https://doi.org/10.17044/scilifelab.20518251.

When comparing different KrakenUniq databases, we found that database size played an important role for robust microbial identification, see Supplementary Information S1. Specifically, small databases tended to have higher false-positive and false-negative rates for two reasons. First, microbes present in a sample whose reference genomes were not included in the KrakenUniq database could obviously not be identified, hence the high rate of false-negatives of smaller databases. Second, microbes in the database that were genetically similar to the ones in a sample, appeared to be more often erroneously identified, which contributed to the high rate of false-positives of smaller databases. For more details see Supplementary Information S1.

Although the technique of filtering the KrakenUniq output by depth and breadth of coverage is reliable for establishing the presence of an organism in a sample, the findings of KrakenUniq have to be authenticated, i.e. their ancient status needs to be confirmed, which is impossible to do with the taxonomic classification approach alone. In other words, KrakenUniq has a fairly low detection error (see Introduction) but cannot control the authentication error because it cannot provide any information about the ancient status of the detected microbes. Furthermore, additional validation in terms of evenness of coverage is a highly desirable step as KrakenUniq filters are currently based on hard thresholds in aMeta and thus may not always be optimal.

To validate the results from the KrakenUniq pre-screening step and further eliminate potential false-positive microbial identifications, aMeta performs an alignment with the Lowest Common Ancestor (LCA) algorithm implemented in MALT [20]. Alternatively, aMeta users can also select Bowtie2 for a faster and more memory-efficient analysis but lacking LCA alignments, see Supplementary Information S2. While being more suitable than Bowtie2 for metagenomic profiling, MALT is very resource demanding. In practice, only reference databases of limited size can be afforded when performing analysis with MALT, which might potentially compromise the accuracy of microbial detection. For more details see Supplementary Information S3. In consequence, we aim at linking the unique capacity of KrakenUniq to work with large databases with the advantages of MALT for validation of results via an LCA-alignment. For this purpose, aMeta automatically builds a project-specific MALT database, based on a filtered list of microbial species identified by KrakenUniq. In other words, the combination of microbes across the samples remaining after depth and breadth of coverage filtering of the KrakenUniq outputs is used to build a MALT database which allows the running of LCA-based MALT alignments using realistic computational resources. We found that this design provides two to six times less computer memory (RAM) usage compared to traditional ways of building and using MALT databases, see Supplementary Figure 6.

Thus, the analysis strategy applied in the aMeta workflow is two-step. First, we pre-screen and classify microbial organisms in aDNA samples with KrakenUniq against the full NT or microbial NT database; a step that can be performed virtually on any computer, even a laptop. Second, we validate the findings by performing MALT LCA-based alignments against a project-specific database comprising microbial species identified at the first step by KrakenUniq. This two-step strategy provides a good balance between sensitivity and specificity of both microbial detection and authentication in aDNA metagenomic samples without imposing a large computational resource burden. On one hand, the KrakenUniq step optimizes the sensitivity of microbial detection by using a large database that would otherwise be technically impossible for MALT to build. On the other hand, the MALT step optimizes the specificity of microbial detection and authentication by performing LCA-based alignments suitable for computing various quality metrics. Note that the two-step design of aMeta minimizes potential conflicts between classification (KrakenUniq) and alignment (MALT) approaches by ensuring consistent use of the reference database. In other words, the MALT database always comprises a subset of microbial reference genomes profiled by KrakenUniq.

As previously emphasized, microbial organisms identified by KrakenUniq and MALT in metagenomic samples need to be checked for their ancient status, i.e. authentication analysis is needed in order to discriminate truly ancient organisms from modern contaminants. For authentication of microbial organisms found in metagenomic aDNA samples, we applied the MaltExtract tool [23] to the LCA-based alignments produced by MALT, and computed the deamination pattern [27, 36], read length distribution, and edit distance (amount of mismatches) [23] metrics. Next, the breadth and evenness of coverage of reads aligned to each microbial reference genome were generated using SAMtools [28]. In addition, the workflow automatically extracts alignments and the corresponding reference genome sequence for each identified microbial organism in each sample, allowing users to visually inspect the alignments, e.g., in the Integrative Genomics Viewer (IGV) [29], which provides intuitive interpretation of the quality metrics reported by aMeta. Finally, histograms of postmortem damage (PMD) are computed using PMDtools [30], which features a unique option of likelihood-based inference of ancient status with a single read resolution. All the mentioned quality metrics are complementary and serve for more informed decisions about presence and ancient status of microorganisms in metagenomics samples. A typical graphical output from aMeta is demonstrated in Figure 4.

**Figure 4.**
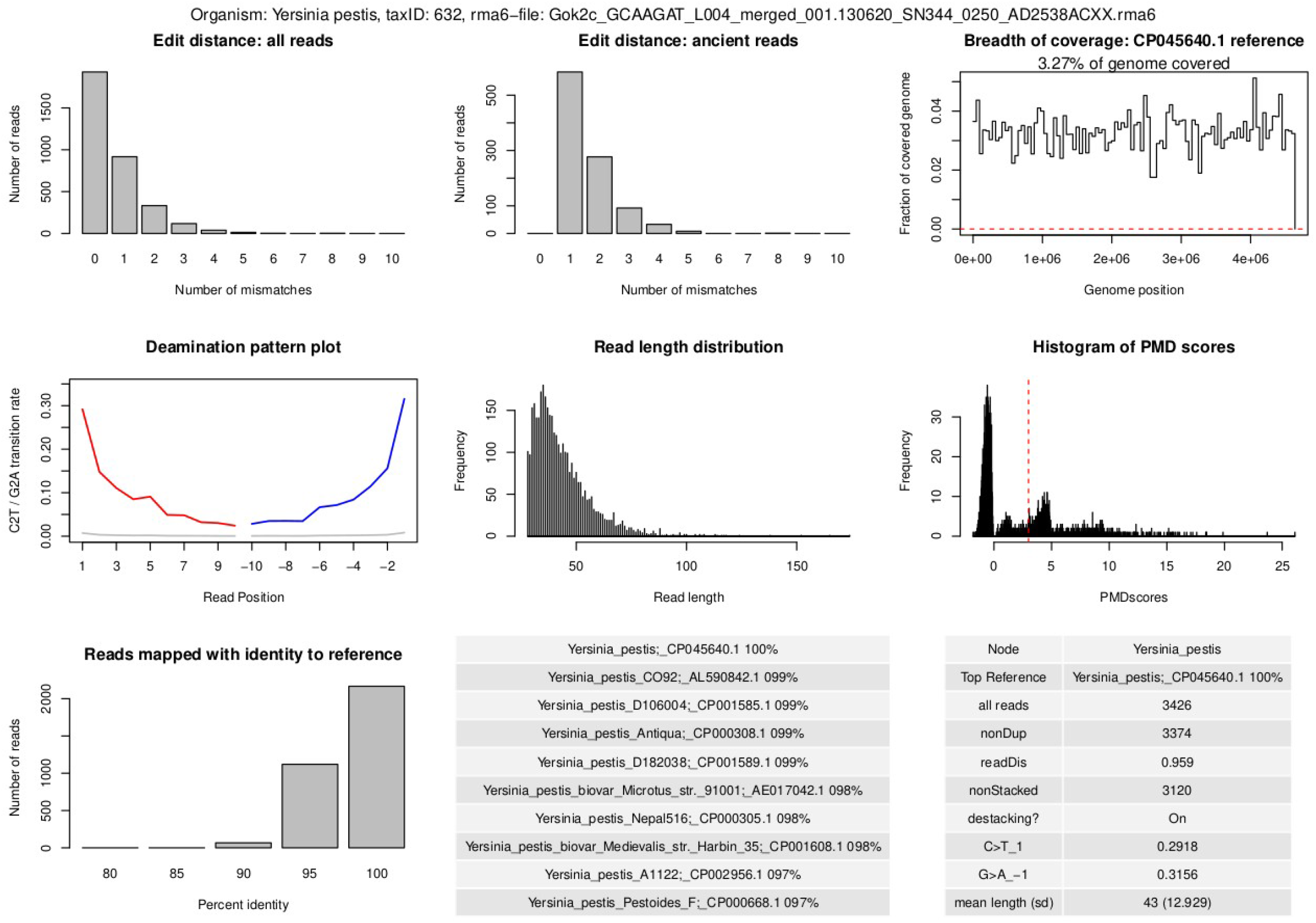
Authentication output of aMeta. Panels from left to right, top to bottom: a) edit distance computed on all assigned reads, b) edit distance computed on damaged reads, c) evenness / breadth of coverage, d) deamination pattern, e) read length distribution, f) PMD scores distribution, g) number of reads assigned with an identity to a reference, h) candidate reference sequences with percentages of mapped reads, i) MaltExtract statistics

In addition to the graphical summary of quality metrics, aMeta delivers a table of microbial abundances quantified from both *rma6*- and SAM-alignments available from MALT. The alignments in *rma6* format are quantified using the *rma2info* wrapper script from the tool MEGAN [31], see Supplementary Information S7, while a custom *awk* script is used for quantifying microbial abundance from SAM-alignments. A disadvantage of *rma6*, which is a primary MALT alignment format, is that it cannot be easily handled by typical bioinformatics software such as SAMtools. However, we found that the alternative alignments in SAM format delivered by MALT lack LCA information and therefore are not optimal either since they essentially resemble the Bowtie2 alignments. Nevertheless, we believe, the two ways of abundance quantification are complementary to each other. The LCA-based quantification from the *rma6* output of MALT might underestimate the true per-species microbial abundance since many short conserved aDNA sequences originating from a species are assigned to higher taxonomic levels, e.g. genus level, and thus do not contribute to the species abundance. In contrast, the LCA-unaware quantification from the SAM output of MALT seems to overestimate the true per-species microbial abundance since it counts absolutely all reads assigned to a species, including the non-specific multi-mapping reads, i.e. the ones that map with the same affinity to multiple homologous microbial organisms.

Within the aMeta workflow we constructed and implemented a special scoring system that should facilitate getting a quick user-friendly overview of likely present ancient microbes. The score is computed per microbe and per sample, and represents a quantity that combines seven validation and authentication metrics presented graphically in Figure 4, more specifically: 1) deamination profile, 2) evenness of coverage, 3) edit distance (amount of mismatches) for all reads, 4) edit distance (amount of mismatches) for damaged reads, 5) read length distribution, 6) PMD scores distribution, 7) number of assigned reads (depth of coverage). The score assigns heavier weights to evenness of coverage as an ultimate criterion for the true presence of a microbe, and deamination profile as the most important evidence of its ancient origin. As one of the outputs, aMeta delivers a list of detected microbial organisms ranked by the score that implicitly corresponds to the joint likelihood of both their presence in a sample and their ancient origin.

### Benchmarking aMeta on simulated data

We benchmarked aMeta against HOPS [23] which is one of the most widely used pipelines in the field of ancient metagenomics. Another popular general purpose aDNA pipeline, nf-core / eager [32], implements HOPS as an ancient microbiome profiling module within the pipeline, therefore we do not specifically compare our workflow with nf-core / eager but concentrate on differences between aMeta and HOPS.

For robust comparison of the two approaches, we built a ground truth dataset which represents 10 ancient human metagenomic samples with various microbial compositions simulated with the tool gargammel [33]. To mimic potential human contamination, we simulated reads that were both endogenous (ancient) and of a contaminant origin (modern). Nevertheless, considerable fractions of DNA reads in each sample were simulated as being of microbial and non-human origin, consistent with microbial contamination, which in our case is of primary interest. We selected 35 microbial species that are commonly found across our aDNA projects [34, 35], and simulated their fragmented and damaged reads. In addition, Illumina adapters and sequencing errors were added to mimic typical ancient DNA raw genomic sequencing data, see Supplementary Information S5 for details. To better resemble a typical situation in our studies [34, 35] where a mixture of various types of microbes is observed, we simulated bacterial reads of both modern and ancient origin. For example, when working with ancient dental calculus [34] one may often observe likely endogenous *Streptococcus pyogenes* or *Parvimonas micra*, which were simulated here as being of ancient origin. In addition to endogenous bacteria, we also simulated a few microbial organisms such as *Mycobacterium avium* and *Ralstonia solanacearum* as ancient, as they can also typically be found in human aDNA samples, while probably being of exogenous, i.e. environmental, origin. The reads of the abovementioned ancient endogenous and exogenous bacteria were simulated to be fragmented and damaged. In total, 18 out of 35 microbial species were simulated as ancient. We also added a number of modern bacterial contaminants such as *Burkholderia mallei* and *Pseudomonas caeni* that were simulated with a moderate fragmentation level and no clear deamination/damage pattern. In total, 17 out of 35 microbial species were simulated as modern. In summary, the simulated ground truth dataset included both human and microbial DNA reads of ancient and modern origin mixed at various ratios with varying levels of damage and fragmentation. We believe that this closely mimics a typical metagenomic composition scenario that we observe in various aDNA metagenomic projects [34, 35].

Using this simulated ground truth dataset, we first sought to quantify the detection error of aMeta and HOPS, i.e., when a tool falsely reports the presence or absence of a microbe in a metagenomic sample, regardless of its ancient status. For this purpose, we ran aMeta on the simulated data using default settings and the full microbial NCBI NT database for the KrakenUniq step. We obtained an abundance matrix of microbial organisms detected by KrakenUniq after filtering for breadth of coverage. For comparison, we also ran HOPS with default configuration parameters using the complete microbial genomes RefSeq database, which was the largest database that was feasible to use for HOPS on a 1 TB of RAM computer cluster node. We quantified the abundance of microbial organisms detected by HOPS using MEGAN [31]. Next, both KrakenUniq and HOPS microbial abundance matrices were filtered using different thresholds for the number of assigned reads, which is equivalent to filtering by depth of coverage. For each depth of coverage threshold applied to the abundance matrices, we compared microbial organisms identified by KrakenUniq and HOPS against the true list of organisms simulated by gargammel. As a criterion of overlap between the prediction and ground truth we used two metrics: Intersection over Union (IoU), aka Jaccard similarity, and F1 score, which both quantify the balance between sensitivity and specificity of microbial detection by KrakenUniq and HOPS, Figure 5. Illustrated by the solid lines in Figure 5, it is demonstrated how IoU and F1 score change at different depth of coverage thresholds applied to the KrakenUniq and HOPS microbial abundance matrices. The dashed horizontal line in Figure 5 corresponds to the IoU and F1 score computed using the depth and breadth of coverage thresholds set by default in aMeta. The default aMeta filtering thresholds were previously empirically determined from the analysis of a number of ancient metagenomic samples [34, 35]. As Figure 5 shows, the default settings of aMeta result in nearly optimal IoU and F1 score values obtained from filtering the KrakenUniq abundance matrix. In Figure 5, one can observe that irrespective of the depth of coverage threshold applied to the KrakenUniq and HOPS abundance matrices, the IoU and F1 score quality metrics for HOPS are always below the sensitivity vs. specificity level provided by KrakenUniq and aMeta.

**Figure 5.**
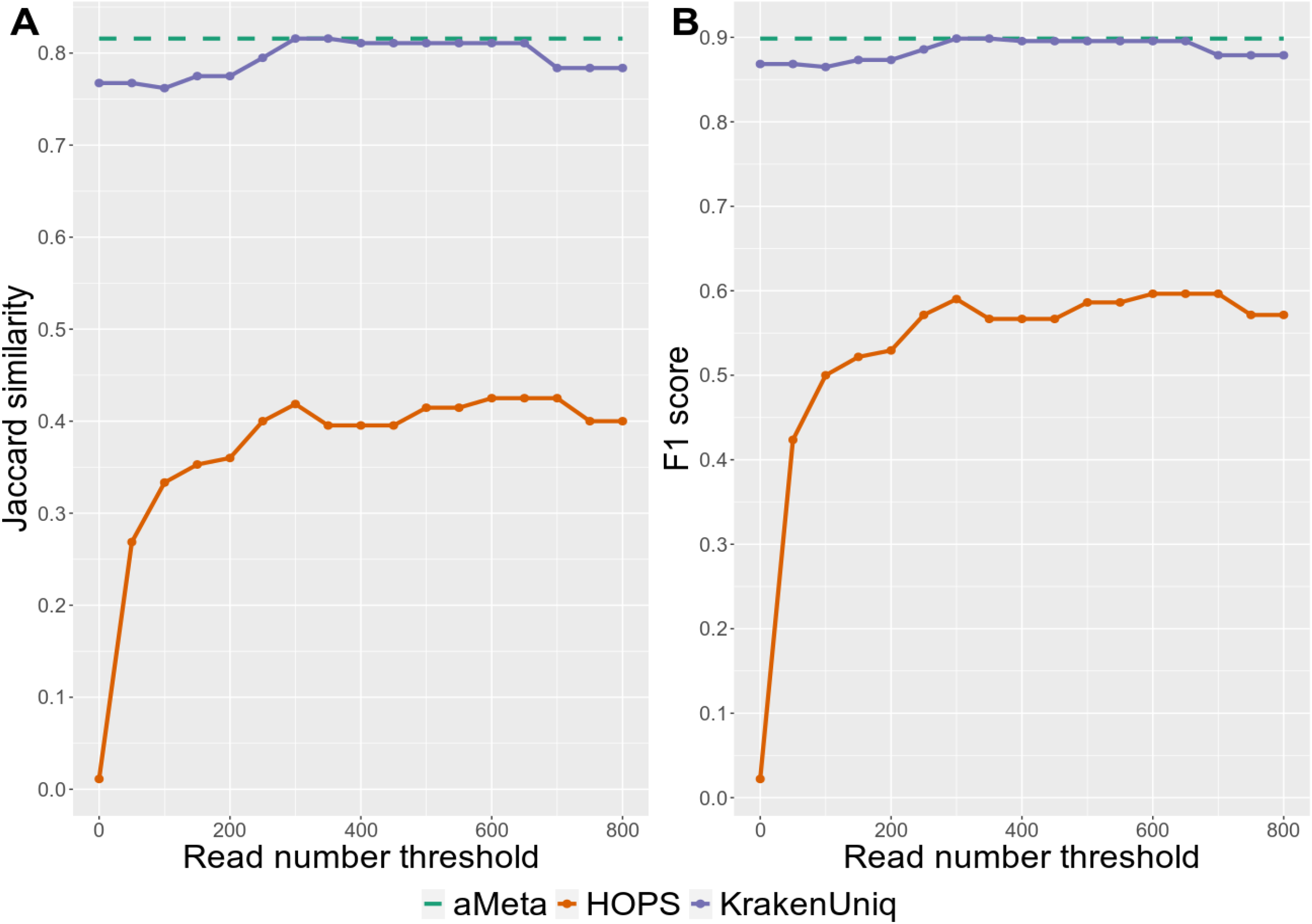
Microbial detection sensitivity vs. specificity comparison between KrakenUniq, HOPS and aMeta at different assigned read thresholds: A) Intersection over Union (Jaccard similarity), and B) F1 score, are computed with respect to the simulated microbial abundance ground truth.

Indeed, this effect comes from two factors. First, since it is computationally feasible to use very large databases for taxonomic profiling with KrakenUniq and hence aMeta, this allows for the detection of microbial organisms that might be missed by HOPS due to their absence in the HOPS database. Therefore KrakenUniq and aMeta have higher sensitivity for microbial detection. This conclusion is confirmed by Supplementary Figures 7–12, where the ground truth for the microbial presence-absence per sample is compared against the one reconstructed by aMeta and HOPS. For example, such simulated species as *Campylobacter rectus*, *Fusarium fujikuroi*, *Methylobacterium bullatum*, *Micromonas commoda, Micromonospora echinospora* and *Mycolicibacterium aurum* were correctly identified by aMeta but not detected by HOPS in any simulated sample. Interestingly, *Campylobacter showae* was detected by HOPS instead of *Campylobacter rectus* because only the former was included in the HOPS database. This shows how a limited database size can impact not only sensitivity (missed microbes) but also specificity (falsely identified microbes) of microbial detection. In total, HOPS missed 12 out of 35 simulated microbial species in all samples, while aMeta missed only 5 out of 35 microbes. The second factor for increased accuracy of microbial detection by aMeta comes from the fact that, while the HOPS microbial abundance matrix can only be filtered based on depth of coverage, an additional breadth of coverage filter is available in KrakenUniq, and hence aMeta, improving the robustness of microbial detection. Therefore KrakenUniq and aMeta tend to have overall higher specificity for microbial detection. For example, *Streptosporangium roseum* was incorrectly identified by HOPS as present in two simulated metagenomic samples, while this species did not pass the breadth of coverage filter applied by aMeta in the two samples, and was correctly excluded from the resulting output. Overall, we conclude that aMeta has a lower detection error compared to HOPS, see Supplementary Figures 7–12 and Supplementary Information S5 for more details.

Further, we addressed the authentication error of aMeta and HOPS, i.e. when a tool wrongly reports a microbe as ancient that was actually not simulated to be ancient. For this purpose, we used the authentication scoring systems implemented in aMeta and HOPS. The scoring systems of both tools not only provide a useful ranking of microbial organisms, but can also be used for computing sensitivity and specificity of microbial validation and authentication for benchmarking purposes. We ran aMeta and HOPS with default settings on the simulated ground truth dataset, and obtained lists of microbial organisms ranked by the scoring system of aMeta and HOPS, where likely present and ancient microbes received higher scores. Visual inspection of the native heatmap output from HOPS demonstrated its poor authentication performance, Supplementary Figure 13. More specifically, a few bacteria such as *Rhodopseudomonas palustris, Rhodococcus hoagii, Lactococcus lactis, Brevibacterium aurantiacum, Burkholderia mallei* were erroneously reported by HOPS to be ancient as they got the highest scores in several samples, while they were supposed to be modern according to the simulation’s design. The native scoring system of HOPS is based on 3 metrics only (edit distance of all and damaged reads + deamination profile), therefore it was carefully generalized to match the scoring system of aMeta, see Supplementary Information S6.

Further, we used the scoring systems of aMeta and HOPS to compute a ROC-curve reflecting sensitivity vs. specificity of microbial validation and authentication by both tools. The comparison of ROC-curves from aMeta against HOPS computed on the gargammel simulated ground truth dataset is presented in Figure 6. One can observe that for the simulated ground truth dataset, aMeta demonstrates overall higher sensitivity vs. specificity of ancient microbial identification compared to HOPS. This is mainly related to the additional evenness of coverage filter and better tuned deamination profile score that helps aMeta establish a more informed decision about microbial presence and ancient status. For example, *Lactococcus lactis* and *Rhodopseudomonas palustris* species which were simulated as modern, obtained high authentication scores from HOPS, which implies that they were predicted to be present and ancient. They were, however, correctly ranked low as potential modern contaminants by aMeta. In contrast, the simulated ancient *Salmonella enterica* genome was ranked low by HOPS due to read misalignment, while it obtained high scores from aMeta correctly indicating its presence and ancient status, see Supplementary Figures 14–15. Overall, we conclude that aMeta has a lower authentication error compared to HOPS, see Supplementary Information S6 for more details.

**Figure 6.**
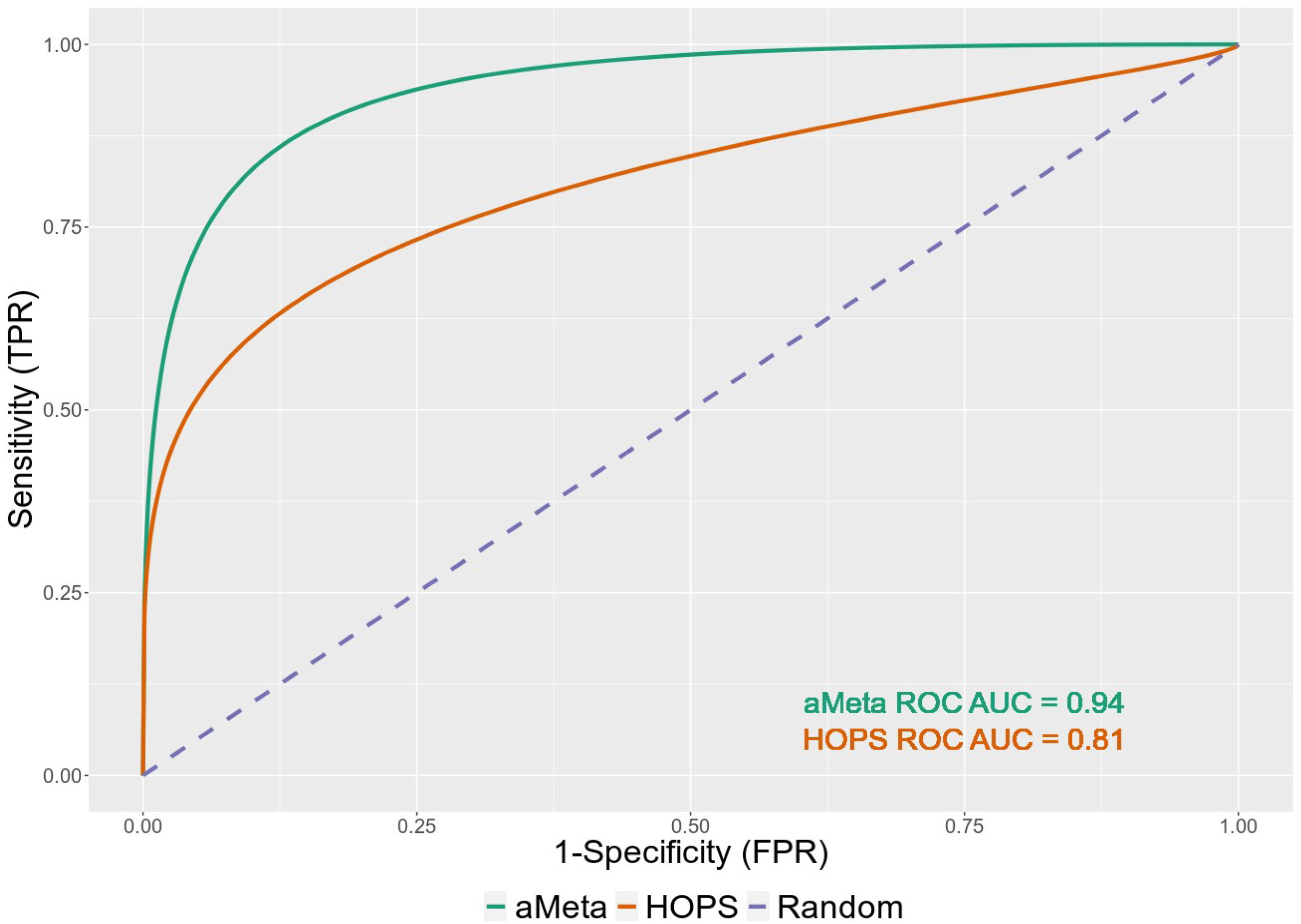
ROC-curve comparison of authentication scoring implemented in aMeta and HOPS.

## Discussion

With the increasing availability of Next Generation Sequencing (NGS) technologies, the field of ancient metagenomics, and particularly of environmental and sedimentary ancient DNA, is currently experiencing rapid growth. Unique archaeological samples such as museum specimens that have been collected and stored for decades can now be analyzed genetically, opening up exciting new opportunities for understanding the past. Technological progress has also been accompanied by method development. While the methodology of traditional ancient genomics reached maturity some time ago, there still does not seem to be a profusion of analytical tools to perform ancient microbiome analysis, presumably because the latter is a much younger field. Currently available ancient metagenomics workflows such as MALT [20], HOPS [23] and nf-core / eager [32] - the latter internally using HOPS - are sensitive to the choice of reference database and are therefore not always optimal in terms of sensitivity vs. specificity balance of microbial detection. In addition, they can be very resource-intensive. Moreover, typical bioinformatics workflows such as nf-core / eager represent rather a collection of tools that provide independent and often inconsistent outputs from taxonomic classification and alignment that need to be manually interpreted and harmonized by users, which is often cumbersome and requires a lot of experience. Therefore, there is currently a need for alternative, more accurate and memory-efficient ancient metagenomics profiling workflows that could be run with minimal user interference.

A challenging peculiarity of ancient metagenomics compared to traditional aDNA analysis is that it involves working with a mixture of various organisms in archaeological samples. In addition, often no clear host is present in a sample, as is the case for environmental and sedimentary aDNA. Therefore, to robustly detect organisms that have left their DNA in archaeological samples, a competitive mapping approach must be used where each query sequence is compared to all reference sequences in a database. The size of a reference database thus becomes an important factor for microbiome profiling as large databases should provide more unbiased identification of present microbes. If the reference database is not large enough, there is a risk, first, of not identifying a microorganism that is not present in the database; and second, of erroneously identifying a microorganism in the database that happens to be phylogenetically close to another microbe truly present in the sample but not included in the database. However, current analytical tools such as MALT [20], HOPS [23] and nf-core / eager [32] can only be run on reference databases of limited size. For more details see Supplementary Information S3. It is therefore important to develop alternative memory-efficient analytical approaches that can screen metagenomic samples against large reference databases.

In this study, we proposed a novel bioinformatics workflow, aMeta, which has a number of advantages over other analytical frameworks in the field. The workflow is based on recent advances in the field of metagenomics and provides a list of ancient microbes robustly detected based on multiple quality metrics with minimal interference from the user. Unlike other typical workflows that often merely combine bioinformatic tools, aMeta was designed to answer a specific research question, which is the robust identification of ancient microbial organisms with optimal sensitivity and specificity of detection and authentication. Therefore, while at first glance our workflow can be seen as a combination of classification via KrakenUniq and LCA-based alignment via MALT, it actually implements a number of additional features that: 1) harmonize the outputs of KrakenUniq and MALT, 2) minimize the amount of manual post-processing work, 3) optimize memory usage, and 4) ensure the user obtains a highly accurate overview of the microbial composition of the query samples.

More specifically, aMeta uses taxonomic pre-screening with KrakenUniq against a large reference database to inform LCA-based alignment analysis with MALT. Initial unbiased pre-screening against large databases becomes computationally feasible thanks to the recent low-memory development of KrakenUniq [22]; meaning, provided that a reference database has been built already and is of a reasonable size, taxonomic classification can be performed on virtually any computer, even a laptop, irrespective of the database size. Indeed, according to our tests, Supplementary Figure 5, the new KrakenUniq development enables 10 times faster classification using a 450 GB reference database even on a computer cluster node with 128 GB of RAM, which was previously impossible without a node with at least 512 GB of RAM. This new development opens up exciting opportunities for truly unbiased pre-screening by KrakenUniq, followed by alignment, validation and authentication by MALT, as implemented in our workflow. In this way, the follow-up with MALT no longer requires superior memory resources; in fact, the memory usage of MALT can be minimized based on the selection of likely present microbial organisms detected by KrakenUniq at the initial pre-screening step. By dynamically building a project-specific MALT database, we ensure that only necessary microbial reference genomes are used for alignment, validation and authentication which dramatically reduces the memory consumption of MALT.

Indeed, our computational memory benchmarking shows that aMeta consumed barely half the RAM compared to HOPS when processing 10 simulated ancient metagenomic samples, Supplementary Figure 6. The memory gain can be explained by two factors. First, despite a larger database used by aMeta (full microbial NCBI NT + human + complete eukaryotic genomes, sequences occupy ~300 GB of disk space) than by HOPS (complete microbial genomes from NCBI RefSeq database, sequences occupy ~60 GB of disk space), the recent fast and low-memory development of KrakenUniq [22] was able to handle the larger database more efficiently and use less memory compared to MALT, which is the implicit engine of HOPS. Second, as a result of pre-screening with KrakenUniq, the dynamically built MALT database had a reduced size compared to the MALT database used for HOPS. In other words, the MALT step in aMeta is not a screening *per se* but a followup after KrakenUniq pre-screening and thus can be performed using a reduced database, unlike HOPS, which is a screening pipeline by design, where in order to obtain an unbiased microbial detection, one has to use a large MALT database which slows down the alignment process as indicated in the Supplementary Figure 6.

Importantly, the memory gain of our workflow does not compromise the accuracy of microbial detection and authentication. Instead, as shown in Figures 5 and 6, aMeta has a better sensitivity vs. specificity balance for both microbial detection and authentication compared to HOPS. On one hand, the superior sensitivity of aMeta comes from a larger reference database used by KrakenUniq compared to the one used by HOPS. In essence, including more microbial organisms into the reference database enables their discovery in query samples. On the other hand, the superior specificity of aMeta is primarily due to robust filtering based on the evenness of coverage applied to candidate microbes. In other words, aMeta does not only rely on the number of reads mapped to a reference genome of a microbial candidate, as does essentially HOPS, but considers the spread of aligned reads across the reference genome as an ultimate criterion of microbial presence. While the evenness of coverage is a crucial metric, aMeta also generates a few other quality metrics such as deamination pattern, edit distance, PMD scores, read length distribution, depth of coverage, and combines them into a score that can be used to rank microbial candidates to get a robust overview of the ancient microbiome.

We believe that the features of aMeta listed above make this workflow stand out in terms of accuracy and resource usage compared to other alternative analytical frameworks in the field. In addition, the Snakemake [24] implementation of the aMeta workflow facilitates reproducibility of the analysis and allows for seamless interaction with high performance (HPC) computer cluster environments, see Supplementary Information S4.

### Limitations and planned extensions of aMeta

The aMeta workflow uses a reference-based approach for the discovery of microbial organisms in metagenomics samples. This implies that only organisms included into a reference database can be found in a sample. Therefore, a current disadvantage of aMeta is that it is not able to discover unknown microbial organisms for which there is no reference genome generated yet.

An alternative approach widely used in modern metagenomics is the *de novo* assembly of microbial contigs. In this case, no prior information about potential microbial candidates is required, and reference genomes can be virtually reconstructed for any microbe present in a sample. This process however typically requires high coverage, i.e. deep sequenced samples, which might be problematic for palaeogenetics where usually a very limited amount of ancient DNA can be extracted from archaeological artifacts. Another complication comes from the ancient DNA damage [36] that, in addition to sequencing errors, complicates the *de novo* assembly process and can lead to the formation of chimeric contigs [37] which could greatly influence the downstream analysis.

A *de novo* assembly module (not presented in this article) written in Snakemake is currently being tested in our lab, and we plan to add it to the workflow in a future release of aMeta. In this way, aMeta will leverage all the power of classification, alignment and *de novo* assembly that can be used complementary to each other and provide a more informative overview of microbial composition in ancient metagenomics samples.

Another planned extension of the aMeta workflow is a special mode for working with ancient environmental and sedimentary DNA, an area of palaeogenetics that has experienced a rapid growth [38]. One challenge here to overcome is the fine-tuning of aMeta workflow for dealing with large eukaryotic reference genomes such as plant and animal genomes. For this purpose, using the non-redundant NCBI NT database may not be optimal as it contains eukaryotic reference genomes that are typically of poor quality and far from complete. Our preliminary testing shows that the large variation in quality of reference genomes across eukaryotic organisms in the NCBI NT database can lead to severe biases in taxonomic assignment of metagenomic reads, where spurious taxa can be detected merely because the taxa have better quality (more complete) reference genomes compared to homologous taxa that are in fact present in the sample.

Further, although the internal default filters used by aMeta are well tuned and seem to demonstrate good performance for the vast majority of aDNA samples [34, 35], we are working on developing a strategy for self-adjusting the filters depending on the nature and quality of aDNA samples. For example, viral organisms have typically small reference genomes, and hence, very few aDNA reads aligned to them. Therefore, hard filtering thresholds that are currently implemented in aMeta might miss rare members of the microbial community and need further tuning which is planned for future versions of the workflow.

Next, although the pre-screening step with KrakenUniq implemented in aMeta substantially reduces the amount of memory needed for performing MALT alignments, we found that large input *fastq*-files (> 500 million sequenced reads) from deeply sequenced samples, or alternatively, a large number (> 1000) of medium-size input *fastq*-files can still result in a severe memory burden for the MALT step that might consume over 1 TB of RAM, even though KrakenUniq is rather insensitive toward the input file size. Therefore, we do not currently recommend merging *fastq*-files from different sequencing libraries corresponding to the same sample as it is often done in genomics projects, but we advise processing *fastq*-files individually unless one has access to very large computer nodes.

Finally, while giving clear advantages compared to HOPS in terms of computer memory usage, aMeta may currently not be as fast as HOPS when extensive multi-threading is available, see Supplementary Figure 6. We, however, are currently working on several optimization schemes that will substantially improve the speed of aMeta in the future release.

## Code and data availability

The workflow is publicly available at https://github.com/NBISweden/aMeta. The non-redundant NCBI NT KrakenUniq database can be accessed at the SciLifeLab Figshare following the address: https://doi.org/10.17044/scilifelab.20205504, and the microbial version of NCBI NT combined with human and complete eukaryotic reference genomes can be accessed via SciLifeLab Figshare at https://doi.org/10.17044/scilifelab.20518251. Further, the Bowtie2 index of NCBI NT is publicly available through SciLifeLab Figshare at https://doi.org/10.17044/scilifelab.21070063, and the pathogenic microbial subset of this index can be access via the SciLifeLab Figshare at https://doi.org/10.17044/scilifelab.21185887. Codes for computer simulations and other scripts used for this article can be accessed at https://github.com/NikolayOskolkov/aMeta.

## Acknowledgements

NO, PU and CM are financially supported by Knut and Alice Wallenberg Foundation as part of the National Bioinformatics Infrastructure Sweden at SciLifeLab.

James A. Fellows Yates, Alexander Herbig, Nico Rascovan, Maxime Borry, Alexander Hübner, Irina M. Velsko, Alina Hiss, Gunnar Neumann and Christina Warinner are greatly acknowledged for providing valuable feedback on the design and technical details of the workflow.

The computations were enabled by resources provided by the Swedish National Infrastructure for Computing (SNIC), partially funded by the Swedish Research Council through grant agreement no. 2018-05973, in particular projects: SNIC 2021/5-335, SNIC 2021/6-260, SNIC 2022/5-100, SNIC 2022/6-46, SNIC 2022/22-507, SNIC 2022/23-275, and Mersin University BAP project 2019-3-AP3-3729. We thank Åke Sandgren at SNIC for his assistance with cluster implementation aspects, which was made possible through application support provided by SNIC.

## Supplementary Information

### S1. Effect of database size

To perform BLAST [26] alignment against the non-redundant NCBI NT database was the gold standard in the early days of metagenomic analysis. Currently, more efficient software such as the Kraken family of tools [16, 17, 21, 22] runs much faster on the NCBI NT. Nevertheless, using a complete NCBI NT non-redundant nucleotide database can still be memory demanding for Kraken tools, therefore we aimed at investigating to what extent one can reduce the size of the NT database without compromising the accuracy of microbial detection. For this purpose, we used a few databases varying in size, and compared the outcomes of KrakenUniq classification on the different size databases, applied to a simulated ancient metagenomic dataset with a known microbial composition, i.e. ground truth, see Supplementary Information S5 for details of the simulations.

We found a strong effect of KrakenUniq database size on robustness of microbial detection, see Supplementary Figure 2. We aimed at reconstructing the ground truth microbial composition known from the simulations by first performing KrakenUniq profiling of the simulated samples and then filtering the results by breadth and depth of coverage. Next, we computed the Jaccard similarity (intersection over union) between the species detected by KrakenUniq in each database and the ground truth species. We used in total four databases varying in size and content. The smallest database used was the NCBI RefSeq complete microbial genomes that included approximately 20 000 reference sequences that together comprised nearly 70 billion nucleotide characters. The largest database was the complete non-redundant NCBI NT that comprised approximately 230 billion nucleotide characters. The intermediate size databases included the Standard Kraken database (default for both Kraken1 [16] and Kraken2 [17]), and the microbial subset of the full NCBI NT, i.e. Microbial NCBI NT.

The smallest NCBI RefSeq complete genomes database gave the lowest Jaccard similarity just below 0.2, which suggests that this database suffers from low sensitivity of microbial detection which may potentially bias taxonomic profiling of metagenomic samples. We found that increasing the database size resulted in gradual growth of Jaccard similarity, i.e. a better overlap of detected species with the ground truth microbial species. Starting with the Microbial NCBI NT database comprising approximately 110 billion characters, the Jaccard similarity reached a plateau at around 0.75. Although the complete NCBI NT, that included both prokaryotic and eukaryotic reference genomes, was able to further increase the Jaccard similarity metric, the effect was rather marginal. This database, however, demanded substantially greater RAM resources. We concluded that the Microbial NCBI NT provides sufficient accuracy when performing microbial profiling, i.e. including eukaryotic organisms into the database (as it is the case for the Full NCBI NT) does not significantly affect the accuracy of microbial detection. Despite the large variation (large error bars) of Jaccard similarity in Supplementary Figure 2, the increasing profile of Jaccard similarity as a function of database size growth is quite clear. Therefore, we concluded that, in our simulation work, the larger databases provide higher accuracy of microbial detection, while smaller databases suffer from low sensitivity and may introduce biases into microbial identification in metagenomic samples.

Further, to demonstrate how spurious mis-alignments may arise when working with small reference databases, we used a random metagenomic stool sample *G69146_pe_1.fastq.gz* from a modern infant from the DIABIMMUNE metagenomic dataset, the Three Country Cohort [39], https://diabimmune.broadinstitute.org/diabimmune/three-country-cohort/resources/metagenomic-sequence-data, and aligned it to *Yersinia pestis (Y. pestis)* CO92 reference genome alone, https://www.ncbi.nlm.nih.gov/genome/153?genome_assembly_id=299265. We discovered that nearly 22 000 reads mapped uniquely, i.e. with mapping quality MAPQ > 0, Supplementary Figure 3. Since the sample was from a modern infant who unlikely suffered from plague, the mapped reads cannot be used as evidence of *Y. pestis* presence in the infant’s stool sample. Further, visually inspecting the alignments in Integrative Genomics Viewer (IGV) [29] we confirmed that the reads aligned unevenly implying *Y. pestis* was not a right reference genome for the reads, see Supplementary Figure 1. Assuming that a large fraction of the aligned reads might be of human rather than bacterial origin, and thus misaligned to the *Y. pestis* reference genome due to the absence of a human reference genome in the reference database, we concatenated the hg38 human reference genome, https://hgdownload.soe.ucsc.edu/goldenPath/hg38/bigZips/, with the *Y. pestis* reference genome and proceeded with competitive mapping. We found, however, that adding the human reference genome to the database did not change the number of reads mapped uniquely to the *Y. pestis* reference genome. Next, we assumed that the ~22 000 misaligned reads originated from microbial organisms (other than *Y. pestis*) that were phylogenetically closer to *Y. pestis* than to humans. We therefore used sequentially: a) 10 random bacterial reference genomes from the NCBI RefSeq database, b) 100 random bacterial reference genomes, c) 1000 random bacterial reference genomes, d) 10 000 random bacterial reference genomes, and finally e) all 28 898 bacterial genomes available from NCBI RefSeq for September 2022, concatenated them with *Y. pestis* + hg38, and performed alignments with Bowtie2 to this concatenated reference. We observed a gradual decrease in the number of reads aligned uniquely to *Y. pestis*: from ~8500 reads at 10 random bacteria down to only 11 reads at 28 898 bacteria, see Supplementary Figure 3. This was a substantial decrease compared to the initial ~22 000 reads, nevertheless we still had a few misaligned reads while our expectation was to observe near-zero reads aligning uniquely to the *Y. pestis* reference genome. We believe that ~10 aligned reads can be treated as a noise level, and therefore should not be considered as evidence of a microbe present in a metagenomic sample. Thus, the increase in database size (the number of reference genomes in the database) allowed us to correctly confirm that *Y. pestis* was not present in the modern infant stool sample.

Overall, we conclude that the database size plays a major role in the robustness of microbial identification in metagenomic samples. A sufficiently small database, while computationally easier to process, may jeopardize the accuracy of metagenomic analysis and lead to high false-positive and false-negative rates for the detection of microbial species.

### S2. Validation of KrakenUniq results via alignment with Bowtie2

Taxonomic classification with KrakenUniq delivers a reliable list of organisms present in a sample. However, the detected organisms have to be validated and their ancient status has to be confirmed. Since this is not possible to do with taxonomic classification alone, additional alignment tracking is needed as it can provide information on coverage and damage of detected organisms. Therefore, aMeta implements the alignments of aDNA sequences to their respective reference genomes using Bowtie2 [19] and MALT [20] aligners. The two aligners are different in terms of mapping methods as well as speed and resource demands. While MALT uses the Lowest Common Ancestor (LCA) algorithm that is preferred for metagenomic data, Bowtie2 is a general purpose aligner that does not assign LCA to each aDNA sequence. Nevertheless, we found that an advantage of Bowtie2 is that it was faster and less resource-intensive than MALT. This has enabled the seamless build and use of large Bowtie2 indexes for the full NCBI non-redundant NT database (currently requires 600 GB of RAM), which is publicly available for the scientific community via the SciLifeLab Figshare at https://doi.org/10.17044/scilifelab.21070063, as well as for a microbial pathogen-enriched database (“PathoGenome”, requires 256 GB of RAM, it is available at the SciLifeLab Figshare via https://doi.org/10.17044/scilifelab.21185887) for fast validation and authentication of potentially present pathogenic species, Figure 1. The latter was built using a custom permissive list of pathogenic microbial organisms derived from the literature. Thus, Bowtie2’s global (end-to-end) alignments can be directly used to compute evenness of coverage, edit distance [23], deamination profile with e.g. mapDamage [27] and other validation and authentication metrics. However, Bowtie2 provides very conservative alignments by not applying LCA. This leads to ignoring aDNA sequences that actually originate from a certain species but cannot be unambiguously attributed to that species due to their length and conservation. These aDNA sequences are marked as multi-mappers by Bowtie2 and are usually removed from downstream analyses. Therefore an additional LCA-based alignment step with MALT [20] is strongly recommended to enhance the robustness of microbiome reconstruction.

### S3. Limitations of MALT and HOPS

The HOPS pipeline [23] was originally developed for screening ancient metagenomics samples to detect the presence of ancient pathogenic microorganisms via LCA-based alignment against a reference database. However, a truly unbiased microbe / pathogen detection is only possible with a sufficiently large reference database. As discussed in Supplementary Information S1, if the size of the reference database is not satisfactory, first, microbial organisms truly present in a sample but not included into the reference database may be missed; and second, reads from microbial organisms not included in the reference database may potentially be misaligned to other phylogenetically related species in the database. In other words, the selection of microbial organisms to include in the reference database may severely bias the outcomes of the screening workflow. We found that MALT and HOPS can be prone to this bias due to their inherently limited ability to use large reference databases. Building a database and running HOPS on even a limited number of RefSeq reference genomes, such as complete genomes, can easily require up to 1 TB of computer memory (RAM), imposing severe limitations on users without access to computer clusters with large memory nodes.

Another substantial constraint of HOPS is that it does not provide information on the breadth and evenness of coverage. As discussed in the main text and shown in Figure 2, this often results in erroneous detection of microbial organisms that happen to have a large number of reads aligned to conserved regions of their reference genomes, see Supplementary Figure 1.

Below, we list a few other minor disadvantages of HOPS that can nonetheless severely impact the analysis outcome if the users are not properly informed. First, vanilla HOPS lacks an adapter trimming procedure, so we experienced that a naive usage of HOPS on raw data can lead to erroneous results when very few reads get assigned to reference sequences. However, running HOPS via nf-core / eager takes care of this issue and should be preferred over the original HOPS. Second, both MALT and HOPS work primarily with the *rma6* alignment format which is non-standard in bioinformatics and cannot be easily handled. This leads to a strong need to use MaltExtract, an extension of MALT, to process *rma6* files. However, MaltExtract is not easily customizable and offers a limited set of operations by default. In particular, it accepts scientific names of organisms instead of taxIDs, which often results in failure to handle an organism in question due to altered annotations in the NCBI databases or due to lack of a valid match between the scientific name and taxID. Moreover, MaltExtract can only authenticate a priori known candidates, for example microbial pathogens, i.e. it does not test all detected microbes. Third, HOPS does not provide alignments in a more traditional SAM format although MALT does. However, the SAM-alignments delivered by MALT lack an LCA implementation and in that sense are equivalent to Bowtie2 alignments, nevertheless they cannot be considered as a good replacement for the Bowtie2 alignments since they in turn lack important quality metrics such as MAPQ. In summary, an issue with using MALT / HOPS alignments is that the *rma6* format with LCA is not very bioinformatic-friendly and cannot be processed with standard bioinformatics tools such as SAMtools; on the other hand, SAM-alignments are either not delivered at all (HOPS), or, if delivered, lack LCA and fundamental alignment quality metrics (MALT). Finally, the HOPS scoring system, more specifically, the *postprocessing.AMPS.r* script belonging to the HOPS toolbox, by default assigns +1 to any microbial hit with non-zero terminal damage. Therefore, even a tiny transition mutation frequency, such as 0.001 at the terminal ends of the reads, counts as the presence of damage, which substantially inflates the authentication error, see Supplementary Figure 13.

Taking into account the shortcomings of MALT and HOPS, such as large resource demands, limited database size, difficulties with *rma6* output format and limitations with filtering by MAPQ and breadth / evenness of coverage, a new generation of ancient metagenomic specific LCA-based aligners is required. A promising alternative that we suggest for future development is to perform alignments with Bowtie2, which is fast and does not need much computer memory (RAM), and to fine-tune the multi-mapping reads with the recently developed sam2lca algorithm [40]. In our opinion, Bowtie2 + sam2lca should entirely replace MALT while delivering alignments in bioinformatics-friendly SAM-format. Other useful metrics provided by HOPS such as deamination pattern and edit distance can easily be computed by mapDamage [27], and by retrieving NM-tag information from the SAM-alignments delivered by Bowtie2.

### S4. Snakemake implementation of aMeta

We implemented the pipeline with the Snakemake workflow management system [24]. The workflow together with installation instructions, documentation and test data set is available at https://github.com/NBISweden/aMeta. The following section gives a brief overview of the implementation, configuration and execution details of the Snakemake workflow.

We followed the best practice guidelines [41] that structure the workflow repository according to the Snakemake workflow template [42] that organizes workflow-related files in a *workflow* directory, and configuration files in a *config* directory. Briefly, workflow command lines, so-called rules, are placed in a *workflow / rules* directory arranged by topics. The files consist of Snakemake codes that define operations on inputs to generate outputs, where the workflow manager determines dependencies between the rules. The modular rules files are connected via a *workflow / Snakefile*, which serves as the main entry point for the workflow. Custom scripts that are executed via rules are included in *workflow / scripts*. In order to enable reproducibility, each rule defines isolated software environments that are deployed with the *conda* package manager [43]. *Conda* environment files are stored in *workflow / envs*. The workflow configuration also supports the use of environment modules commonly used in high performance cluster systems HPC.

In addition to the workflow execution files described above, the directory *workflow / schemas* stores configuration *schemas* that define sample data and configuration file formats. *Schema* validation is applied to ensure the validity of sample sheets and configuration files. The configuration file mainly defines sample-sheet location and database resources. An example of a configuration file and a sample sheet can be found in the GitHub repository of the workflow at https://github.com/NBISweden/aMeta.

The general advice for running the workflow is to clone the repository and organize data files following the directory structure guidelines. Then, given a configuration file *config / config.yaml* and sample sheet, the workflow can be executed from the root directory of the repository. Compute resource usage, such as memory and run time, can be further fine-tuned on a rule-by-rule basis through the use of *snakemake profiles* [44].

As mentioned above, the execution order of different Snakemake rules is determined by the workflow manager creating a Directed Acyclic Graph (DAG) of jobs that can be automatically parallelized. A typical DAG of aMeta run is presented in Supplementary Figure 4.

### S5. Simulating ancient microbial sequencing data

We used the tool gargammel [33] to simulate 10 metagenomic samples with varying human and microbial composition. Both endogenous (ancient) and contaminant (modern) human reads were present in all the simulated samples. In total, 35 microbial species (31 bacteria, 2 amoeba, 1 fungus and 1 algae) commonly found in our ancient metagenomic projects [34, 35], were simulated with varying abundance across the samples. The abundance of each microbe in a metagenomic sample was set randomly, however the fractions of human and microbial organisms were made to sum up to 1. We simulated reads belonging to 18 ancient and 17 modern microbes. The list of simulated microbial organisms is shown below:

#### Ancient

Campylobacter rectus, Clostridium botulinum, Enterococcus faecalis, Fusarium fujikuroi, Mycobacterium avium, Mycolicibacterium aurum, Neisseria meningitidis, Nocardia brasiliensis, Parvimonas micra, Prosthecobacter vanneervenii, Ralstonia solanacearum, Rothia dentocariosa, Salmonella enterica, Sorangium cellulosum, Streptococcus pyogenes, Streptosporangium roseum, Yersinia pestis, Bradyrhizobium erythrophlei

#### Modern

Acanthamoeba castellanii, Aspergillus flavus, Brevibacterium aurantiacum, Burkholderia mallei, Lactococcus lactis, Methylobacterium bullatum, Micromonas commoda, Micromonospora echinospora, Nonomuraea gerenzanensis, Pseudomonas caeni, Pseudomonas psychrophila, Pseudomonas thivervalensis, Vermamoeba vermiformis, Rhodococcus hoagii, Rhodopseudomonas palustris, Mycobacterium riyadhense, Planobispora rosea

For ancient microbial reads, we implemented deamination / damage pattern with the following Briggs parameters [27, 36] in gargammel: *-damage 0.03,0.4,0.01,0.3*. The simulated ancient reads were fragmented and followed a log-normal distribution with the following parameters *--loc 3.7424069808 --scale 0.2795148843*, that were empirically determined from the *Y. pestis* reads in another project [34]. Illumina sequencing errors were added with the ART module of gargammel to both modern and ancient reads. Finally, Illumina universal sequencing adapters were used, which resulted in 125 bp long paired-end reads. Each simulated metagenomic sample contained 500 000 ancient and modern DNA fragments. The codes used for generating ground truth microbial abundances as well as simulating ancient metagenomic reads are available on GitHub: https://github.com/NikolayOskolkov/aMeta.

Supplementary Figure 7 demonstrates a ground truth microbial abundance heatmap across the 10 simulated samples. The elements of the matrix correspond to the simulated fraction of a microbe in a metagenomic sample. The ground truth abundance matrix was binarized using 1% of all reads as a detection threshold, and a binary (present / absent) heatmap of distribution of microbial organisms across samples is presented in Supplementary Figure 8. To compare the microbial detection accuracy of HOPS and aMeta, we built binary heatmaps based on the microbial abundances reconstructed by HOPS and aMeta, see Supplementary Figures 9 and 10, and compared them with the ground truth in Supplementary Figure 8. For aMeta, default internal filter settings were used to determine the presence / absence of a microbe, while we used a reasonable threshold of 300 reads to binarize the HOPS microbial abundance matrix. Nevertheless the exact threshold value does not significantly affect the conclusions as we show later. From Supplementary Figures 8, 9 and 10, one can conclude that both HOPS and aMeta failed to reconstruct the following simulated species: *Aspergillus flavus, Planobispora rosea, Prosthecobacter vanneervenii, Pseudomonas caeni* and *Vermamoeba vermiformis*. These species were either absent from the reference databases or their abundance was below the detection thresholds in all samples.. However, it is clear that the sensitivity of HOPS was worse compared to aMeta since, in addition to the above mentioned species, HOPS missed *Acanthamoeba castellanii, Bradyrhizobium erythrophlei*, *Campylobacter rectus*, *Fusarium fujikuroi*, *Methylobacterium bullatum*, *Micromonas commoda, Micromonospora echinospora*, *Mycobacterium riyadhense, Mycolicibacterium aurum*, *Nonomuraea gerenzanensis*, *Pseudomonas psychrophila*, *Pseudomonas thivervalensis* in all 10 simulated samples while aMeta mostly detected the species correctly in some of the samples. In total, HOPS missed 12 out of 35 microbial species in all samples, while aMeta missed only 5 out of 35 microbes in all samples. Therefore, undiscovered species contribute to the very high false-negative detection rate of HOPS. In contrast, the specificity of HOPS was comparable to that of aMeta which itself was defined by internal default filters applied to KrakenUniq output. Nevertheless, as an example of non-specific hits, HOPS incorrectly reported *Streptosporangium roseum* in samples 5 and 6 where this species was not actually present, whereas aMeta had more conservative filters that correctly identified this species as not present. A plausible explanation for the misidentification by HOPS is that its less diverse database was unable to correctly disentangle reads originating from *Streptosporangium roseum* and another closely related species, *Planobispora rosea*.

To summarize the results, we computed the accuracy of microbial detection, i.e., presence or absence irrespective of the ancient status, based on the confusion matrices for aMeta and HOPS, see Supplementary Figure 11. The confusion matrix corresponding to aMeta had a detection accuracy of 86% and was more balanced compared to the HOPS confusion matrix, which had an accuracy of 69% and a very high false-negative rate. While a comparable numbers of false-positive discoveries were reported by both HOPS (11 false-positive hits) and aMeta (14 false-positive hits), the false-negative rate of HOPS was nearly three times higher (97 false-negative hits) than that of aMeta (34 false-negative hits), resulting in an overall accuracy of microbial detection by aMeta that was much higher than that of HOPS. It is important to note that the higher accuracy of microbial detection from aMeta was not due to improper filters, i.e. the 300 reads threshold applied to the HOPS microbial abundance matrix, but due to the fact that many species were not present in the HOPS database based on complete microbial NCBI RefSeq genomes. Note that despite that absence, the NCBI RefSeq complete genomes database required almost twice the memory (RAM) resources when executing HOPS, Supplementary Figure 6. To verify the effect of HOPS filtering, we checked the range of different numbers of assigned reads (depth of coverage) thresholds varying from 0 (no filtering) to 800 reads (very harsh filtering), see Supplementary Figure 12. Within the tested range of read number cutoffs, the accuracy of HOPS microbial detection was consistently lower than that of aMeta and did not show substantial variation. This confirms the superior microbial detection accuracy of aMeta.

### S6. Scoring systems of aMeta and HOPS

The scoring system of aMeta represents a sum of seven validation and authentication metrics computed on MALT LCA-based alignments: 1) deamination profile, 2) evenness of coverage, 3) edit distance for all reads, 4) edit distance for damaged reads, 5) read length distribution, 6) PMD scores distribution, and 7) number of assigned reads (depth of coverage). Each metric can add +1 to the total sum except for the evenness of coverage that can add +3, as it is the most crucial to validate microbial presence, and deamination profile, which can add +2 (both 5’ and 3’ ends vote independently), as it is the ultimate criterion for ancient status. Therefore, the range of scores a microbe can obtain varies from a minimum of 0 to a maximum value of 10. Below we explain how each validation and authentication metric presented in Figure 4 is quantified according to the aMeta scoring system.

First, the deamination profile represents an increased frequency of transition mutations (C→T and G→A) at the terminal ends of the sequencing reads compared to the frequencies of all other possible mutations. The scoring system of aMeta adds +1 to the total score if the frequency of C→T exceeds 0.05 at 5’ end and +1 if the frequency of G→A exceeds 0.05 at 3’ end. The exact thresholds can be adjusted depending on the age and preservation conditions of the samples. However we have found that the default thresholds provide sufficient accuracy in most typical cases [34, 35]. Second, the evenness of coverage metric can add +3 to the total sum in the case where the aDNA reads cover the reference genome uniformly. The large contribution of evenness of coverage to the total score is due to its importance for validating the presence of a microbe in a sample. To quantify the evenness of coverage in Figure 4, we split the reference genome into 100 tiles and count the number of tiles with near zero average breadth of coverage (percent of covered bases in a tile). If fewer than 3 tiles have an average breadth of coverage of less than 1%, i.e. if a few aDNA reads are present in almost all tiles, the coverage is considered sufficiently uniform and the total score is increased by 3 points. Note that the mean breadth of coverage across the whole genome may be quite low in this case, for example, only 2-4%. This may reflect the overall low sequencing depth in the case a sample underwent a shallow aDNA sequencing. However, what is most important is that there are reads (even very few) present in nearly all regions of the reference genome. This is what the evenness of coverage metric aims to address. Third and fourth, the edit distance for all reads and damaged reads can add +1 point each, provided that they both have a decreasing profile of the numbers of mapped reads as the number of mismatches increases. This metric controls that the majority of aDNA reads map with no or very few mismatches (damaged reads must have at least one mismatch) by analogy with HOPS [23], and therefore ensures that the reads are mapped to a correct reference genome. We examine the monotonous decline in the numbers of mapped reads by checking that it is always greater for smaller mismatch values. Fifth, the fragmentation of aDNA is inspected via the read length distribution. If 90% of the reads are less than 100bp in length, which ensures that the mode of the read length distribution is located at smaller values typical for aDNA, this quality metric adds +1 to the total score. Sixth, an alternative to the deamination profile assessment of post-mortem damage (PMD) can be performed via PMDtools [30] which computes a likelihood of being ancient for each DNA read. As stated in the original paper [30], a PMD score greater than 3 implies a high likelihood that a read is of ancient origin. This quality metric adds +1 to the total authentication score if at least 10% of the reads have a PMD score above 3. Seventh and last, the depth of coverage is controlled by a 200 DNA reads threshold, which is an empirical number of reads sufficient to compute a statistically reliable deamination profile. If there are more than 200 reads mapped to a reference genome, this adds +1 to the total score.

The native HOPS scoring system includes only three quality metrics: deamination profile and edit distances for all reads and damaged reads. This not only results in many obvious false-positive discoveries, as shown in Supplementary Figure 13, but also too little variation in scores that is not straightforward to align with aMeta’s seven-metric scoring system to compute a ROC-curve. Therefore, for a proper comparison, we had to modify the native HOPS scoring system and add reasonable assessments of depth and evenness of coverage, as well as PMD scores and read length distribution. The deamination profile metric can still, by analogy with aMeta, add a maximum of +2 to the total HOPS score if the transition frequencies at the terminal ends of the aDNA reads are greater than 0, i.e. if any deamination (not necessarily a strong one) is present. Note that this threshold of 0 is designed by HOPS and hard-coded within the *postprocessing.AMPS.r* script belonging to the HOPS pipeline. The edit distances of all reads and damaged reads can also contribute +2 as long as they have declining profiles. Those three native HOPS metrics are not modified by us at all, but only scored accordingly to agree with the scoring system of aMeta. Next, HOPS actually provides read length distribution and depth of coverage metrics but does not use them for its native scoring system. For a more correct comparison with aMeta, we added the read length distribution and depth of coverage metrics to the total HOPS score and quantified them in exactly the same way as within the aMeta scoring system. Further, since the evenness of coverage filter is completely absent from HOPS, it could have been naturally quantified such that HOPS assigns a score of +3 to any microbe, regardless of whether it has uniform coverage or not. However, extending the HOPS scoring system in this manner would obviously worsen its performance. Therefore, based on the correlation between depth and breadth of coverage generally assumed until recently in the ancient metagenomics community for the reason explained in Figure 3, we assigned +3 score if there were more than 200 reads aligned to a microbe by HOPS. Finally, since HOPS does not compute PMD scores, we tried to reasonably add it to the HOPS score by postulating that this quality metric can contribute +1 to the total score if the transition frequency at both terminal ends of the reads exceed 0.05. In summary, the native HOPS scoring system included 3 quality metrics (deamination profile, edit distance of all reads, edit distance of damaged reads) and we left them unchanged. Two other quality metrics (read length distribution and depth of coverage) were straightforward to add since HOPS actually computes them even if it does not use them for scoring, so we added these two quality metrics in exactly the same way as within aMeta. The only two quality metrics that were not provided by HOPS at all, evenness of coverage and PMD score distribution, were added to the total HOPS score in the most reasonable and fair way we could think of. Note that generalizing the native HOPS scoring in order to match the aMeta scoring does not worsen but rather improves its performance over the original HOPS authentication scoring, which is more prone to a high false-positive rate.

We used the aMeta scoring system to predict the ancient status of each microbe in the 10 simulated samples, resulting in 174 predictions as scores varying from 0 to 10. By analogy, we used the modified HOPS scoring for each microbe detected by HOPS and obtained 97 predictions. The different number of predictions between aMeta and HOPS can be explained by a better sensitivity of aMeta described in the main text and Supplementary Information S5. The predictions obtained by aMeta and HOPS were compared to the simulated ground truth ancient / modern labels which allowed computing sensitivity vs. specificity ROC-curves of microbial authentication.

### S7. Other technical details

Below we provide a few unrelated but presumably important technical details about our analysis.

Breadth and evenness of coverage of reads aligned to each microbial organism was addressed using the *samtools depth* command from SAMtools [28].

Sequencing adapters were removed prior to the HOPS run, as this step is not implemented in HOPS by default.

To quantify microbial organisms from the output *rma6*-files of HOPS, the *rma2info* command from MEGAN tool [31] and its wrapper, rma-tabuliser, developed by James A. Fellows Yates [45] were used.

By default we filter output of KrakenUniq using 1000 unique *k*-mers (breadth of coverage filter), and 300 reads (depth of coverage filter) assigned to microbial species. Therefore, the dashed horizontal line in Figure 5 corresponds to the IoU and F1 score computed using the default depth (300 assigned reads) and breadth (1000 unique *k*-mers) of coverage thresholds set in aMeta.

F1 score was computed according to the formula F1 = (2 * TP) / (2 * TP + FP + FN), while the IoU metric, aka Jaccard similarity, was computed as IoU = length of intersection between prediction and ground truth / length of union between prediction and ground truth.

ROC-curve was computed from aMeta and HOPS scores by using *rocit* function from *ROCit* R package https://cran.r-project.org/package=ROCit with *binormal* method.

The evenness of coverage plot in Figure 4 visualizes the results of *samtools depth* command executed on a BAM-alignment. This command outputs the number of reads covering each base of a reference genome. The evenness of coverage is calculated though splitting a reference genome into 100 tiles and computing average breadth of coverage, i.e. percent of covered bases, in each tile.

An additional speed advantage of aMeta comes from optimization of all steps with GNU parallel [46] that is extensively used, for example, for computing deamination profiles with mapDamage [27] in parallel for a number of microbial organisms.

---

**Supplementary Figure 1.**
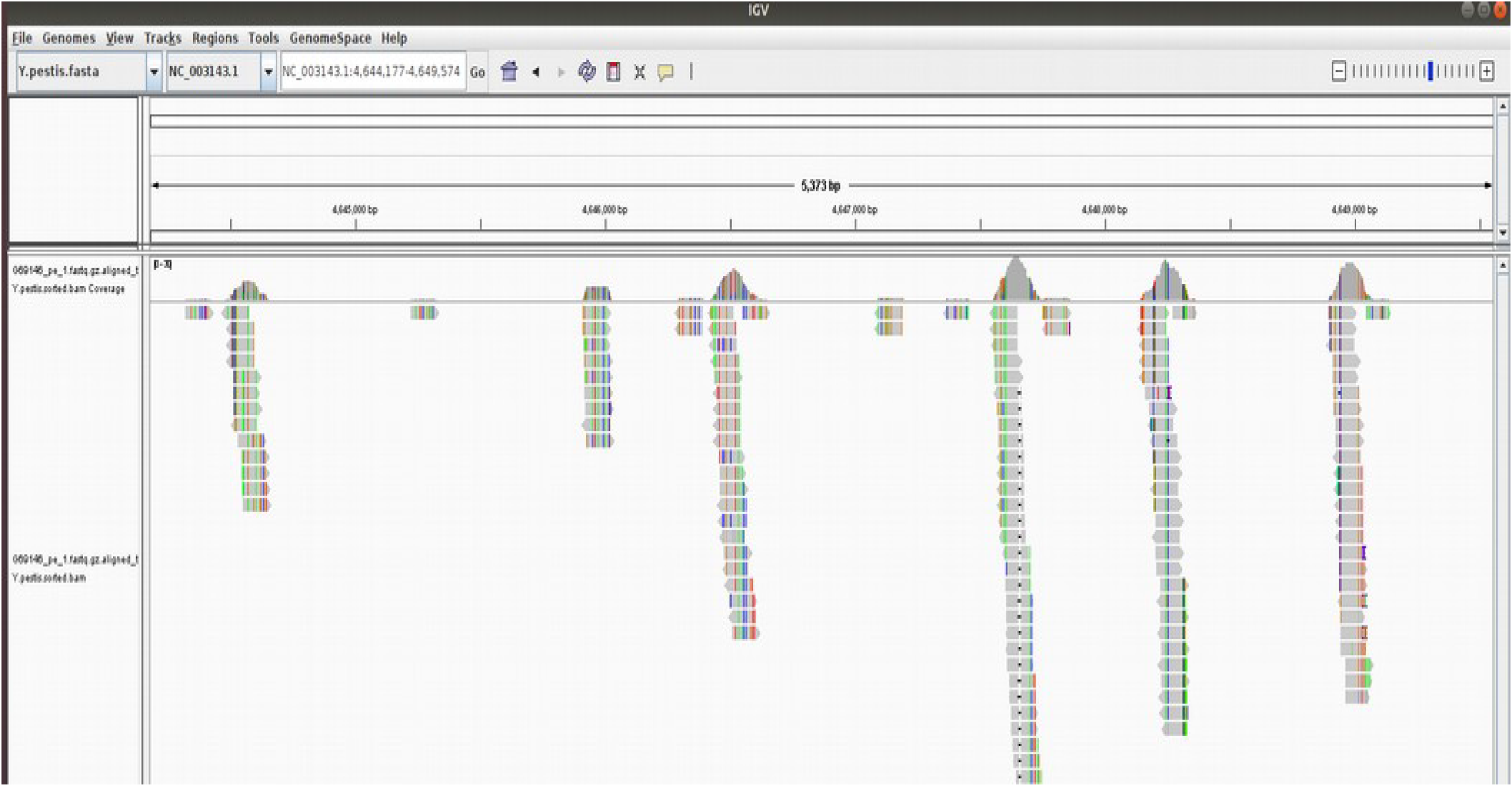
Visualization of false-positive alignments of modern metagenomic reads to *Yersinia pestis* reference genome in IGV. Metagenomic reads from a stool sample taken from a modern infant (who unlikely had a plague) were mapped against a *Y. pestis* CO92 reference genome alone, which resulted in a number of reads aligned in a non-uniform way to regions presumably conserved across bacterial organisms.

**Supplementary Figure 2.**
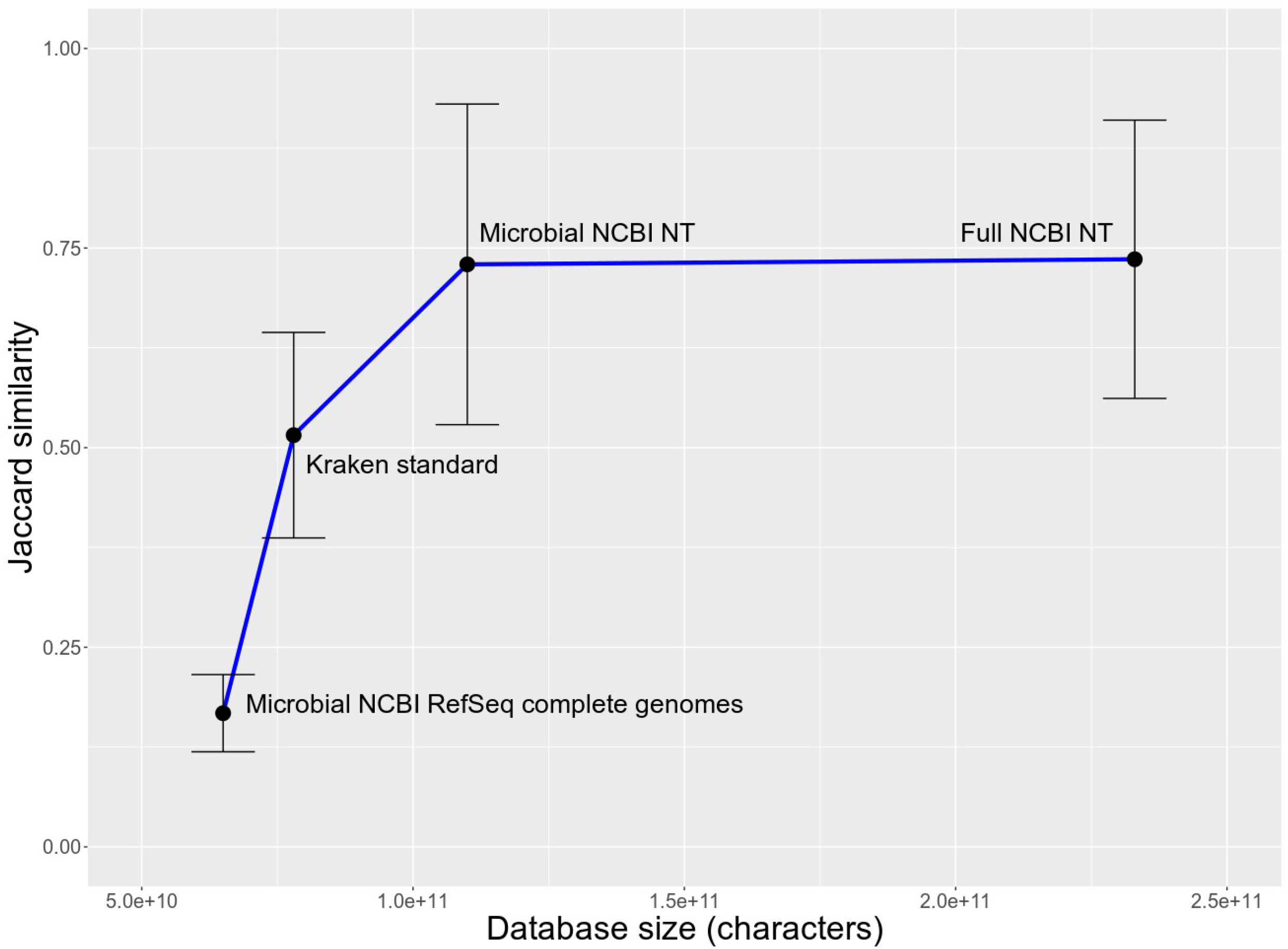
Effect of database size on microbial identification with KrakenUniq: Jaccard similarity (intersection over union) metric was computed with respect to a simulated ground truth. Larger databases tend to provide a better overlap with the ground truth.

**Supplementary Figure 3.**
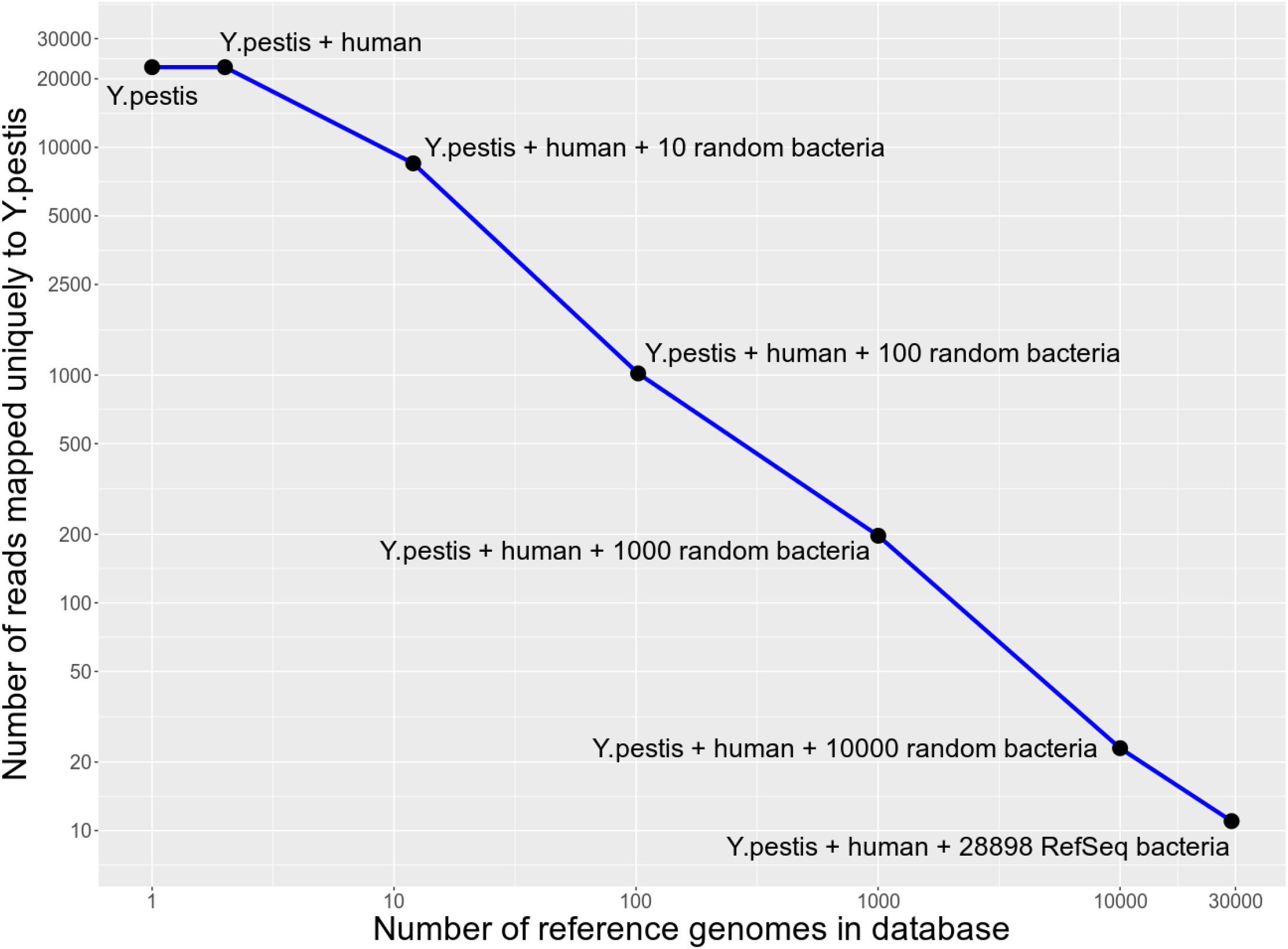
Effect of database size on the number of modern metagenomic reads uniquely mapped to *Yersinia pestis* CO92 reference genome. Starting with *Y. pestis* alone, ~22 000 reads map uniquely. This number gradually decreases down to only a few reads with the growth of the database, i.e. when the human hg38 reference genome is added, followed by adding 10, 100, 1000 and 10 000 random bacteria, and finally all available 28896 bacteria from the NCBI RefSeq database. The axes of the plot are log10-scaled.

**Supplementary Figure 4.**
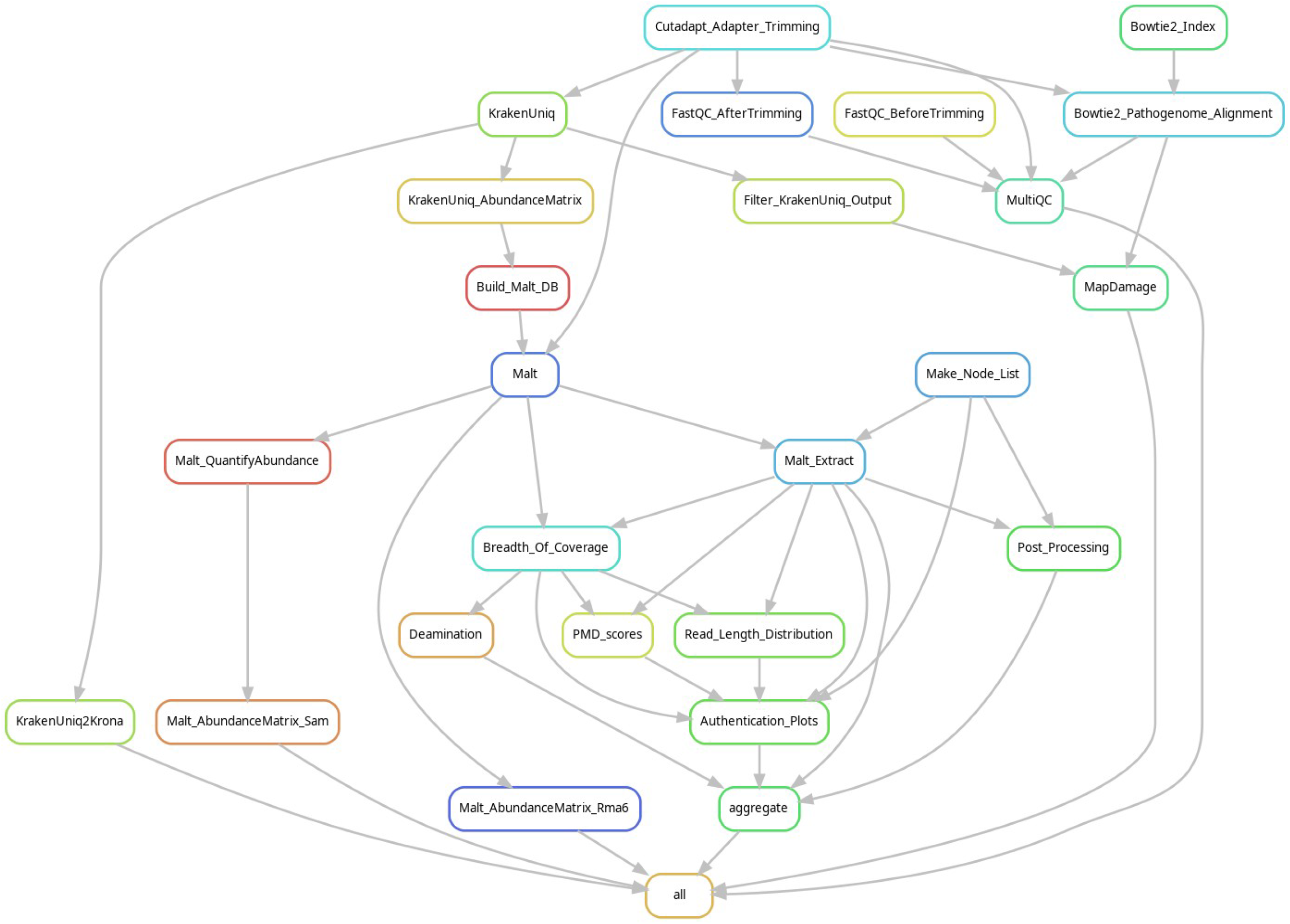
Directed Acyclic Graph (DAG) of a typical project run via the aMeta workflow.

**Supplementary Figure 5.**
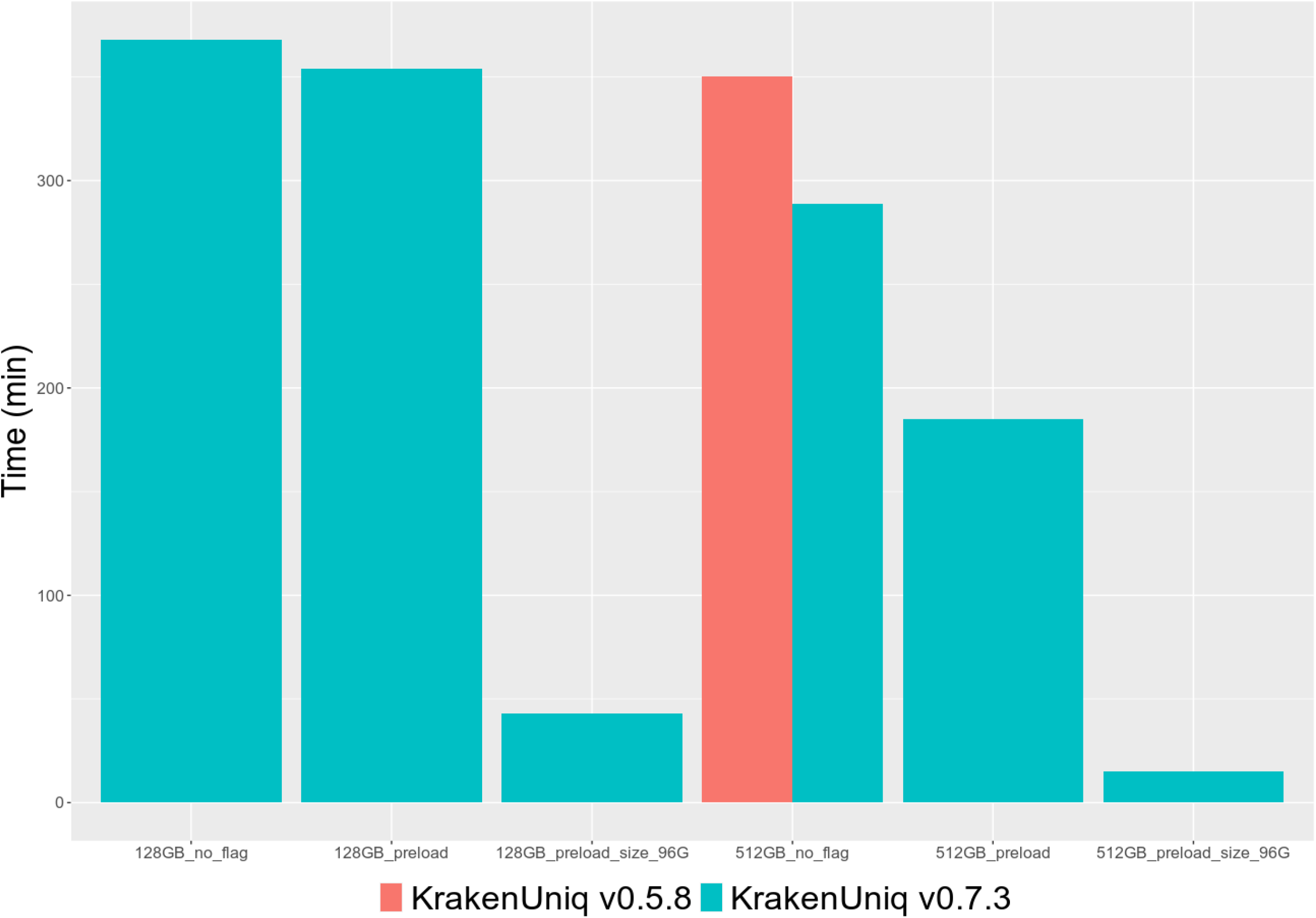
Benchmarking KrakenUniq memory requirements for version 0.5.8 and low-memory version 0.7.3. The database used was ~450 GB in size and could only be fit into a 512 GB memory computer node when using the version 0.5.8. However, database chunking implemented in v0.7.3 allowed processing the same data on 128 GB RAM computer node, and nearly 10 times faster.

**Supplementary Figure 6.**
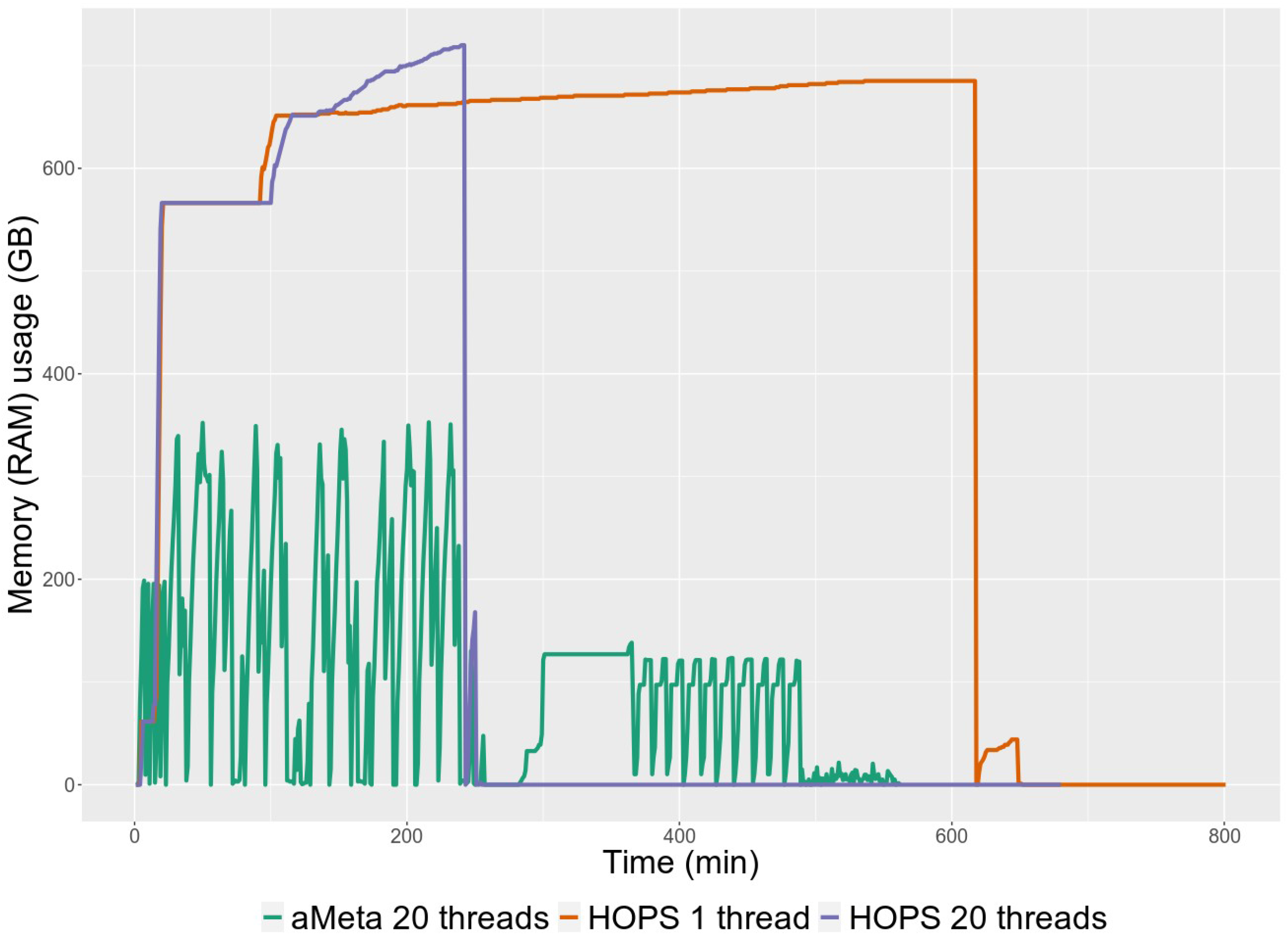
HOPS vs. aMeta memory (RAM) usage comparison.

**Supplementary Figure 7.**
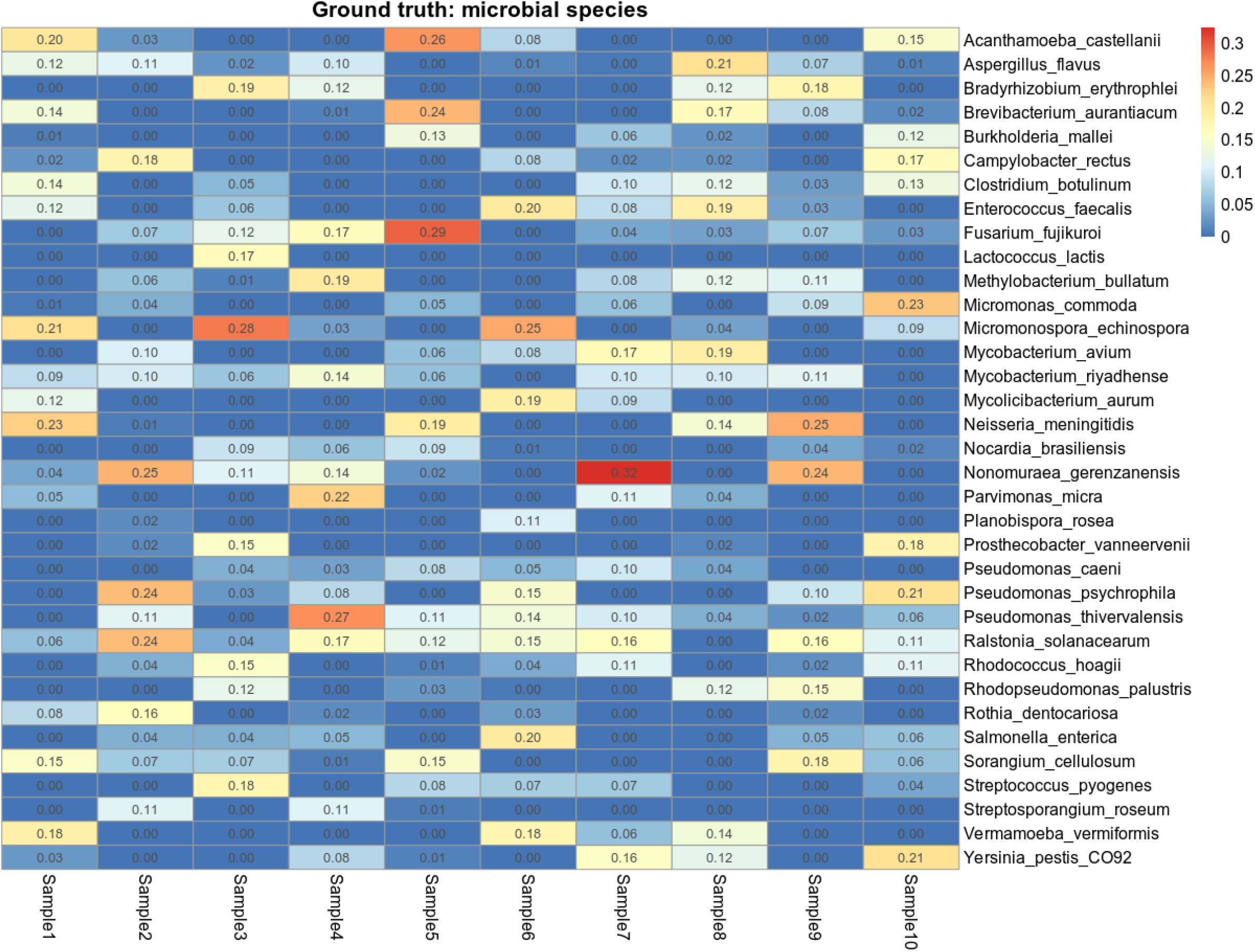
Heatmap demonstrating ground truth microbial abundance in each simulated sample. The elements of the matrix correspond to the simulated fractions of microbes in metagenomic samples (0 – microbe absent, 1 – microbe present).

**Supplementary Figure 8.**
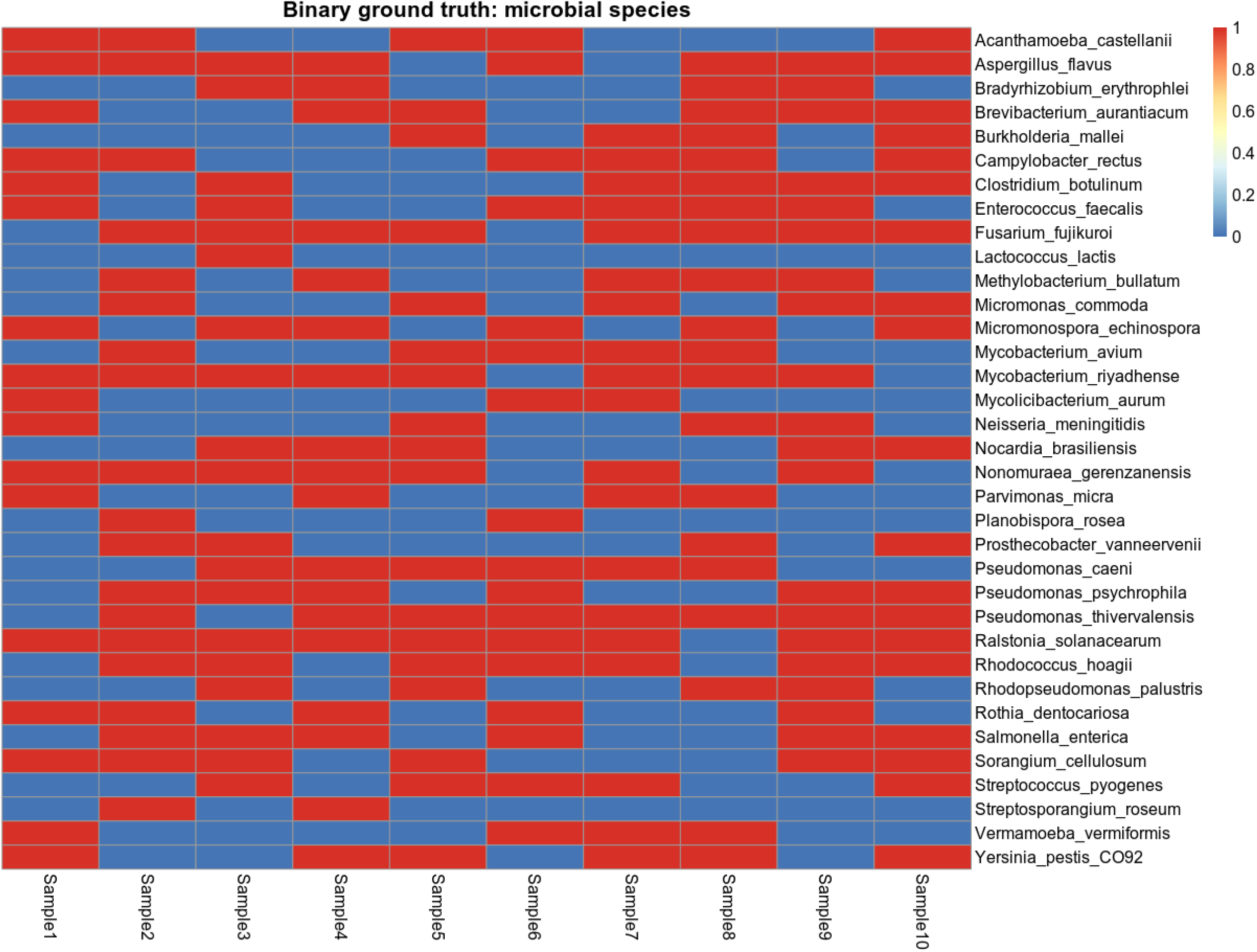
Heatmap of binarized (present / absent) ground truth microbial abundance in each simulated sample.

**Supplementary Figure 9.**
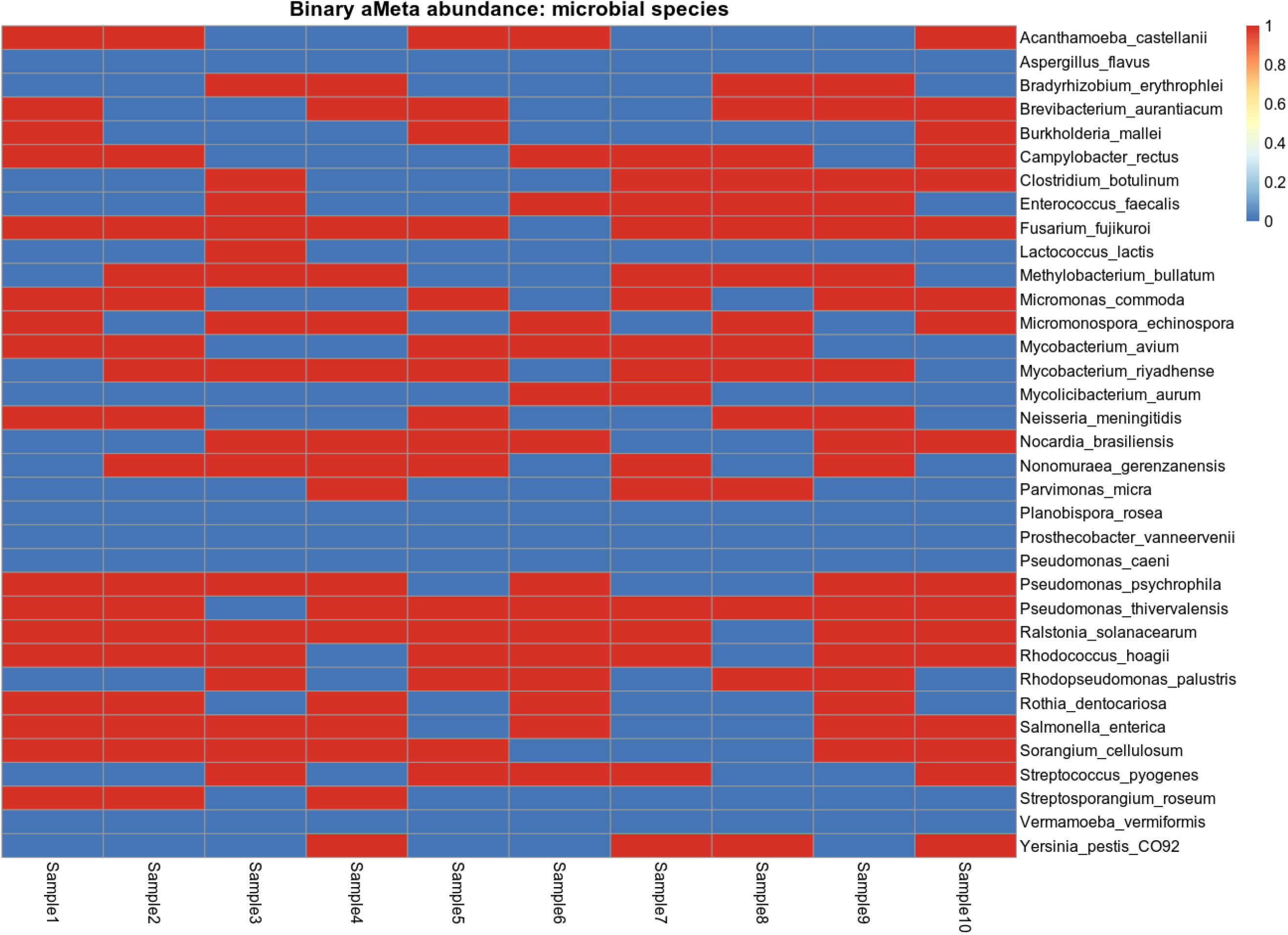
Heatmap demonstrating binarized (present / absent) microbial abundance reconstructed by aMeta.

**Supplementary Figure 10.**
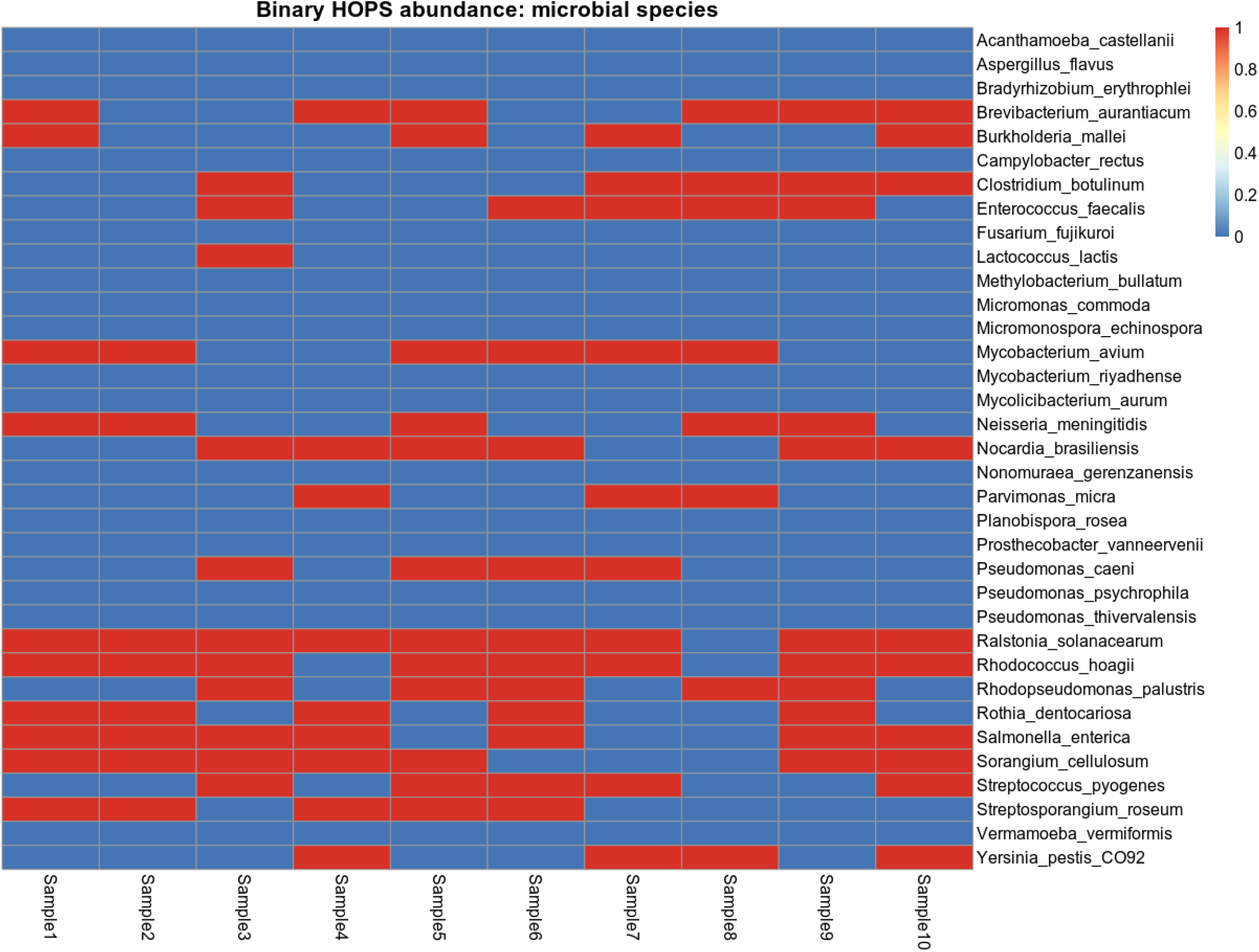
Heatmap demonstrating binarized (present / absent) microbial abundance reconstructed by HOPS.

**Supplementary Figure 11.**
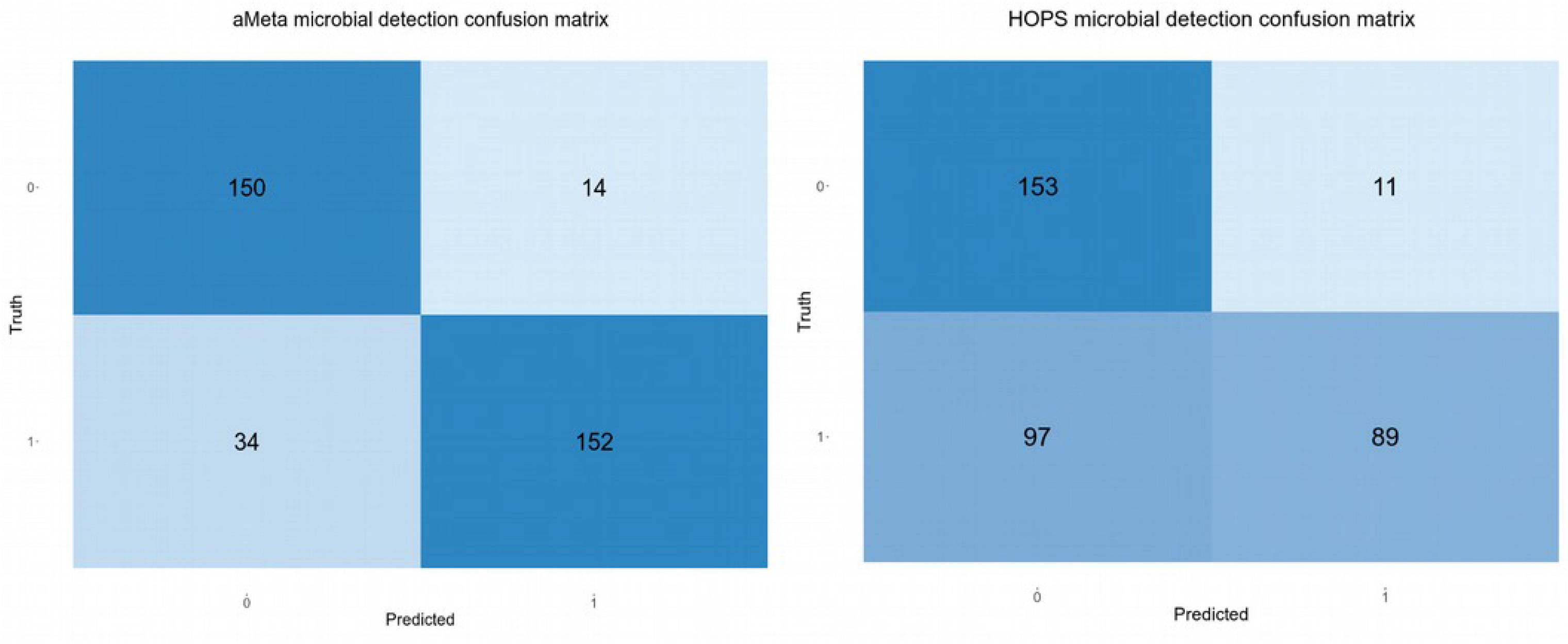
Confusion matrix of microbial reconstruction by aMeta and HOPS: 0 – microbe absent, 1 – microbe present.

**Supplementary Figure 12.**
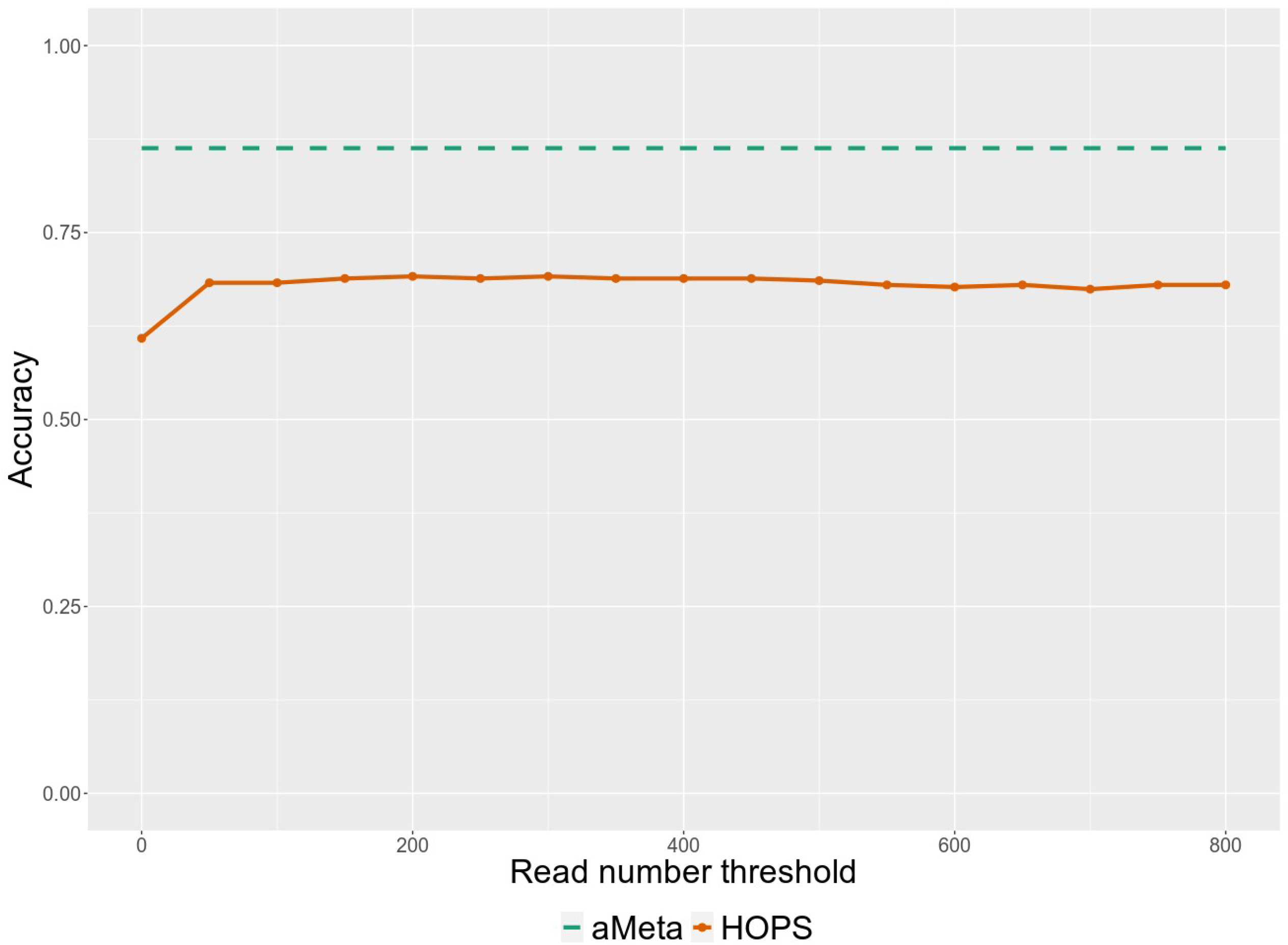
Microbial reconstruction accuracy in simulated samples by aMeta and HOPS.

**Supplementary Figure 13.**
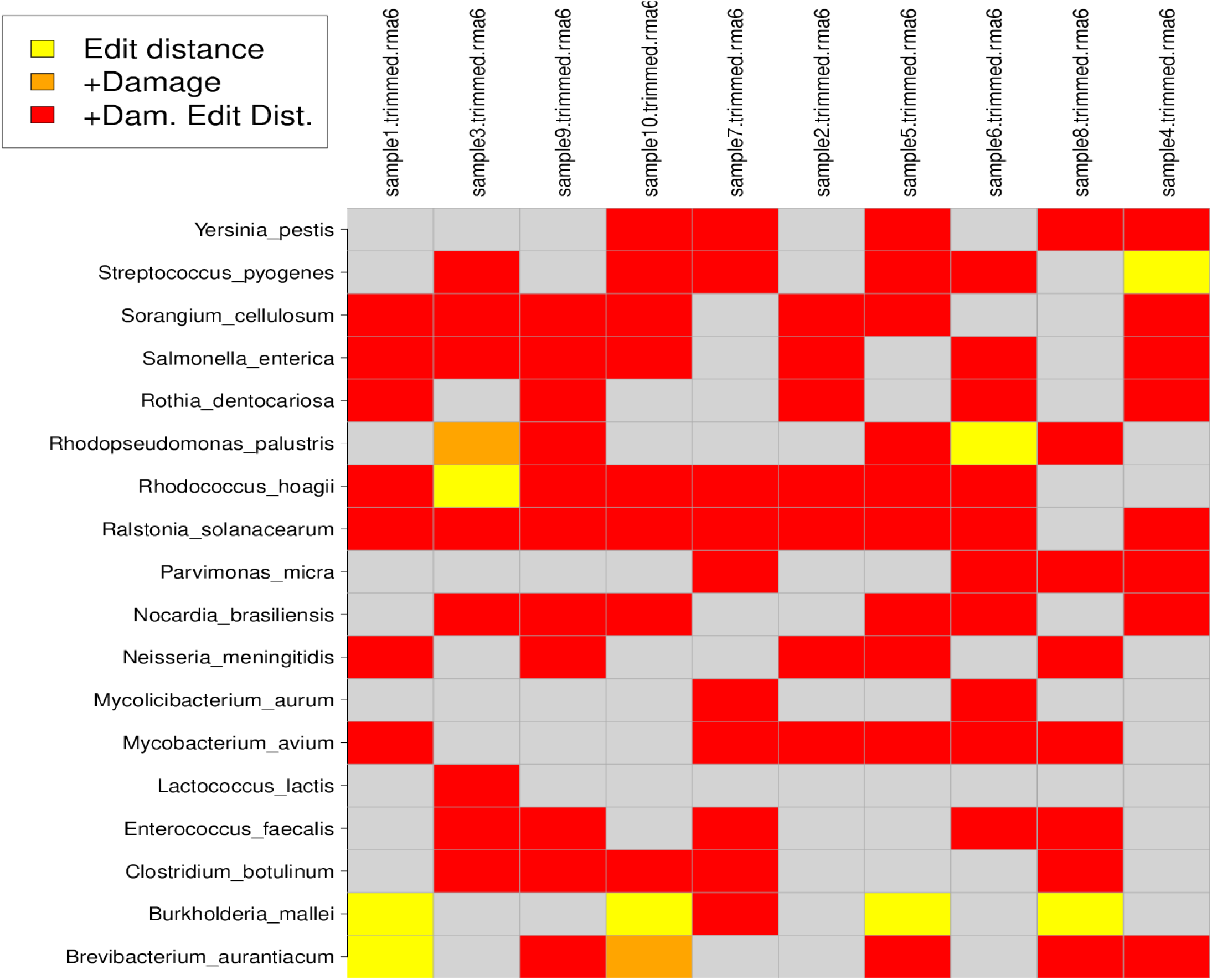
Heatmap of HOPS authentication scores that shows different levels of confidence of ancient microbial presence in simulated data. *Rhodopseudomonas palustris, Rhodococcus hoagii, Lactococcus lactis, Brevibacterium aurantiacum, Burkholderia mallei* were simulated modern, while the other species on the y-axis of the heatmap were simulated ancient.

**Supplementary Figure 14.**
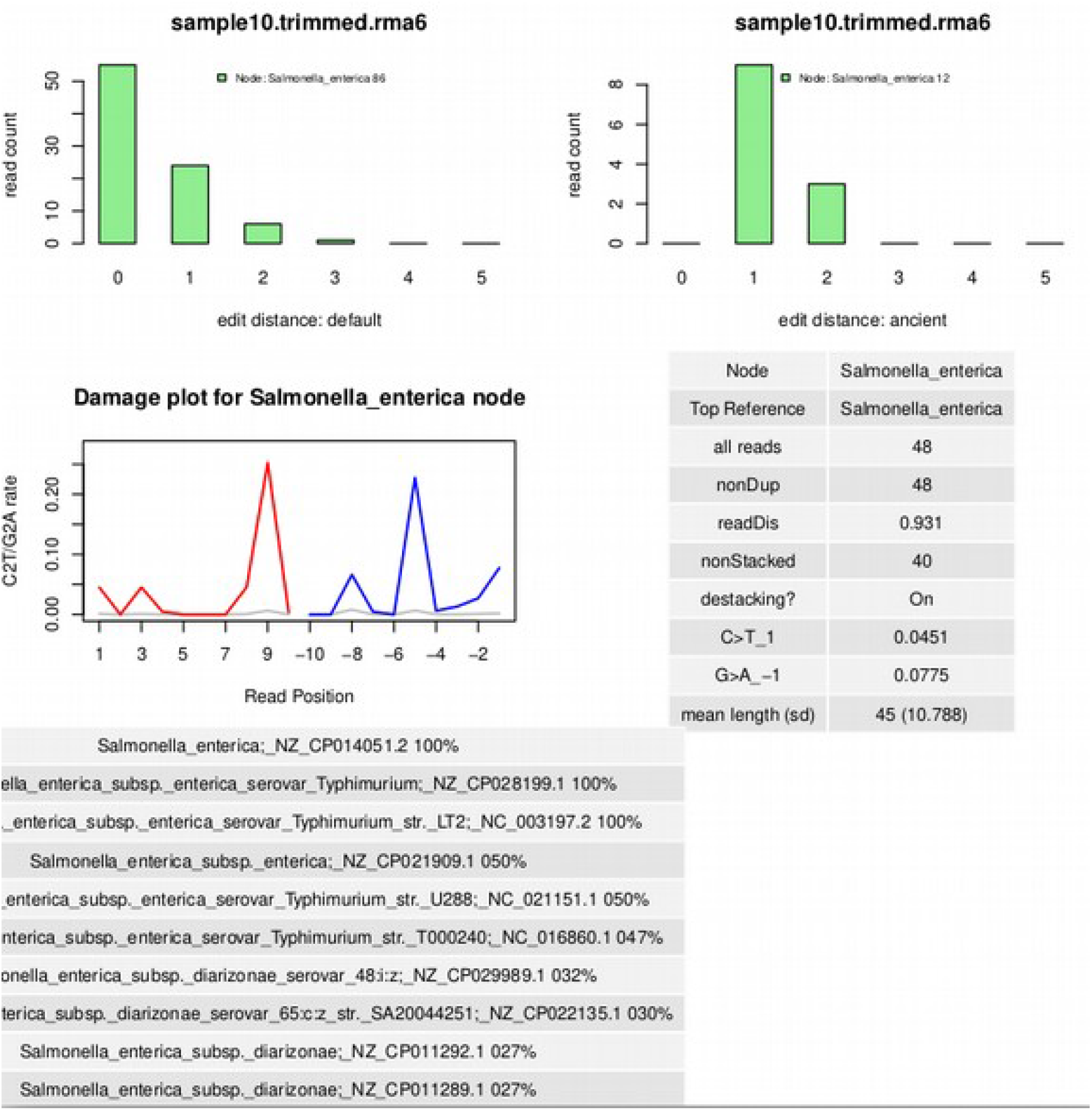
Authentication output from HOPS for *Salmonella enterica* that was simulated to be ancient in sample 10.

**Supplementary Figure 15.**
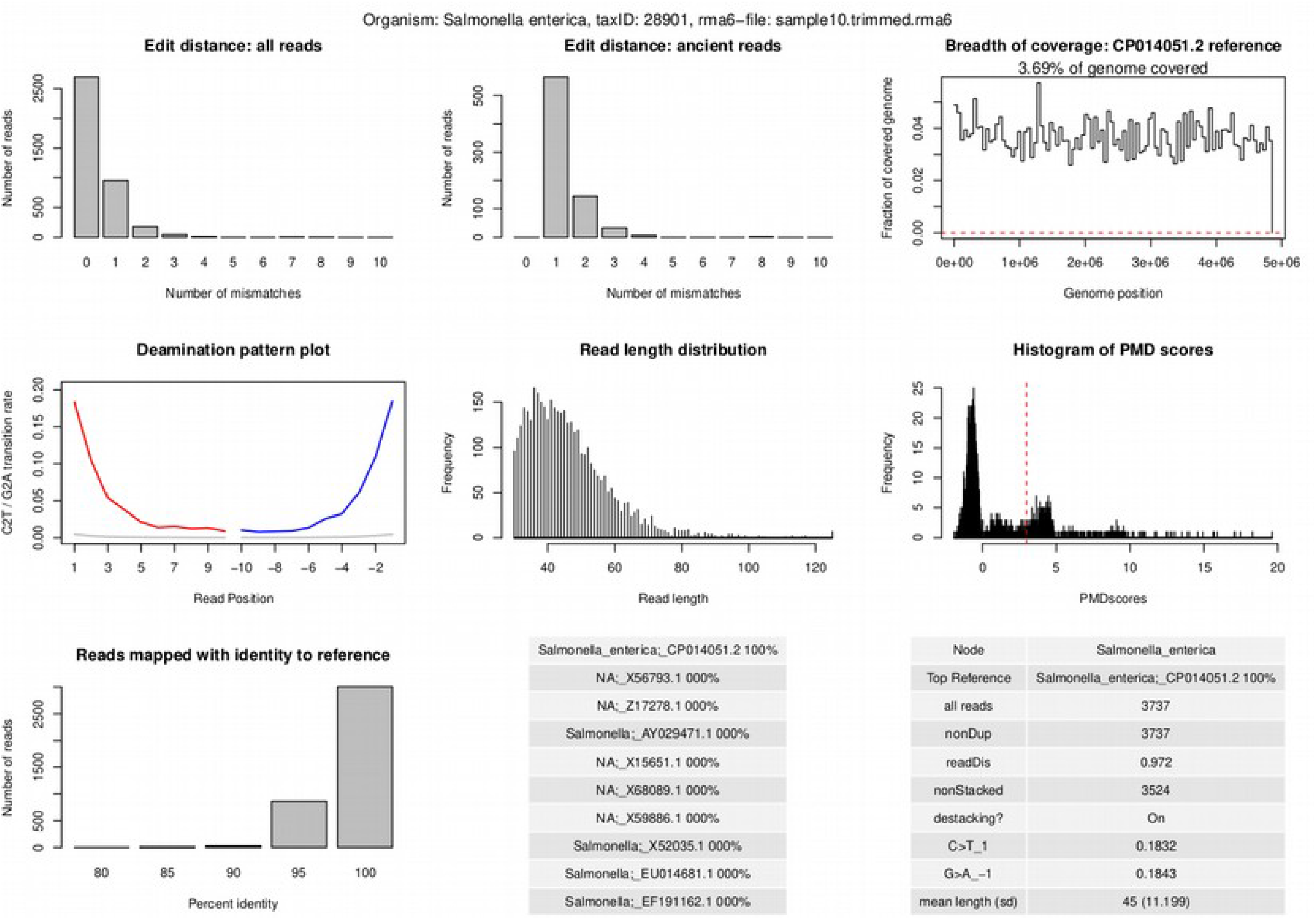
Authentication output from aMeta for *Salmonella enterica* that was simulated to be ancient in sample 10.

